# Octave: Scale-Resolved Evaluation of Spatial Gene Expression Prediction from Histology

**DOI:** 10.64898/2026.09.25.754484

**Authors:** Qing Wang, Sipei Gu, Qizhen Lan, Xiaoqian Jiang, Yijiang Chen, Qianqian Song

**Affiliations:** Department of Health Outcomes and Biomedical Informatics, University of Florida, Gainesville, FL, USA; Department of Health Data Science and Artificial Intelligence, McWilliams School of Biomedical Informatics, UTHealth Houston, Houston, TX, USA; Department of Medicine, College of Medicine, University of Florida, Gainesville, FL, USA; School of Applied and Creative Computing, Purdue University, West Lafayette, IN, USA

**Author notes:** These authors contributed equally (Qing Wang, Sipei Gu).

## Abstract

Spatial gene expression prediction from histology is typically evaluated by the mean per-gene Pearson correlation (PCC). PCC does not distinguish spatial scales: coarse tissue organisation alone can earn most of the score. A domain oracle makes this precise: given an image-derived partition of the tissue, the predictor that assigns each domain its measured mean expression is the best that any predictor constant on that partition can do. On a widely used benchmark with a ∼100 *µ*m pitch, a learnable version that estimates each domain’s mean from training data reaches 82% of the PCC of a trained model, and a partition of the spot coordinates alone, given measured means, reaches 123%. At that pitch the score rewards coarse spatial structure whatever its source. We introduce Octave, a scale-resolved evaluation that filters prediction and measurement into spatial bands with widths in micrometres and scores their agreement in each band. On 16 *µ*m Xenium data, differences that look modest under PCC grow at finer scales: at ∼20 *µ*m the oracle’s shortfall relative to the trained model is 2.4 times its PCC shortfall at the median and exceeds it in all 13 specimens. Across 57 image encoders the finest band reorders 261 of the 1596 encoder pairs relative to PCC. The benchmark’s own finest band is 50 *µ*m wide, coarser than the scale at which these differences appear, so no score on that pitch can directly validate finer structure. Octave reports how well a model predicts spatial expression at each spatial scale.

## 1 Introduction

Predicting spatial gene expression from an H&E image is now a standard benchmark task, with a growing family of methods (He et al., 2020; Zeng et al., 2022; Xie et al., 2023; Chung et al., 2024) competing on a public leaderboard (Jaume et al., 2024). Almost all report the same metric, the mean over genes of the Pearson correlation between predicted and measured expression (Jaume et al., 2024; Mejia et al., 2024; Gindra et al., 2025). The leading methods differ by a few hundredths of that mean.

A leaderboard margin shows that one method scores higher. It does not show what the margin is made of. Tissue is organised into domains such as tumour, stroma and immune infiltrate, which differ enormously in expression and are visually obvious in H&E. A predictor that assigns one expression profile to tumour and another to stroma, with no variation inside either, already correlates well with the measurement; yet the point of measuring expression in space is the structure inside those domains, at the scale of a few cells. The two parts sit at different spatial scales, so the same correlation measured at each scale can separate them, and a scalar correlation cannot.

Our hypothesis is that a predictor allowed only the identity of image-derived domains recovers most of the reported score, and that what it misses is the structure inside them. We make it exact with a *domain oracle*: partition the tissue into clusters of image appearance, each fragmented across the section rather than one connected region, and give every location the measured mean expression of its cluster. Built from the measured expression of the sample it scores, the oracle is not a model but an upper bound on what any predictor constant within those clusters can score; a learnable version takes the cluster means from training data instead. To find where the agreement lives, we introduce Octave: diffuse prediction and measurement on the graph of measurement locations, take the difference between successive smoothing levels to isolate one spatial scale, and correlate the two fields at each scale, with every scale expressed in micrometres. We make three contributions:

1. **Domain identity alone earns most of the score**. The oracle beats a trained model for 56 of 57 frozen encoders on a public benchmark (Jaume et al., 2024), and with means learned from training data the same partition still earns 82% of the model’s score there and 79% on Xenium (Janesick et al., 2023).
2. **Resolved over scale, the gap the scalar hides grows as the scale gets finer**. In the finest band the oracle’s correlation is 0.04 against the trained model’s 0.24; at 20 *µ*m its shortfall is 2.4 times its scalar shortfall, in 13*/*13 specimens; and across fifty-seven encoders the finest band reorders 261 of the 1596 pairs that the scalar ranks.
3. **The standard benchmark cannot reach the scale where the gap appears**. Its spots lie ∼ 100 *µ*m apart, a pitch at which its finest band is 50 *µ*m wide, coarser than the 10–30 *µ*m at which the oracle and the model separate on Xenium. No score on this pitch can directly validate structure finer than that band.

A margin on the reported score therefore does not show that finer structure was recovered (Section 3.2). Where the data resolve that structure, Octave measures its recovery with no new data, baseline or retraining; below the pitch, only finer data can.

## 2 Method

This section makes the hypothesis of Section 1 exact and builds the measure. Section 2.2 bounds what domain identity can score, Section 2.3 shows that giving the score a physical scale returns the score itself, and Section 2.4 defines the readout that does not; Sections 3.2 and 3.3 test them.

### 2.1 Setup

Let **Y** ∈ ℝ^*N ×G*^ hold measured expression for *G* genes at *N* spots with coordinates *x*_1_, …, *x*_*N*_ ∈ ℝ^2^, and let Ŷ be a prediction on the same spots. The standard metric is

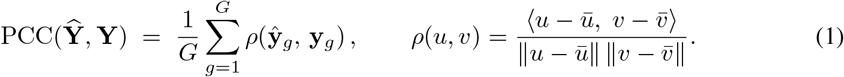

A gene whose prediction or measurement is constant on the scored spots has no correlation; we score it as 0 on Xenium and Visium HD, and the HEST-Benchmark pipeline omits it from the mean.

#### Multiscale operator

Build a symmetric *k*-nearest-neighbour graph on *x*_*i*_ with adjacency **A** and degree **D**, and let

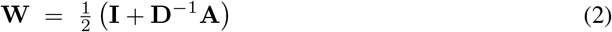

be the lazy random walk. Take diffusion times *t*_0_ = 0 and *t*_*i*_ = 2 ^*i−*1^ for *i* = 1, …, *L* with *L* = 12, and define the low-pass operators 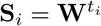.

We measure the physical width of each low-pass operator. Diffuse a unit impulse **e**_*s*_ at spot *s* for *t* steps and take the second moment of the resulting kernel,

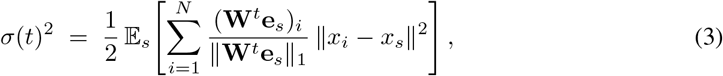

where the factor 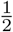 makes *σ* the per-axis width in two dimensions. Over three orders of magnitude *σ* grows as *t*^0.43^ to *t*^0.49^, close to the 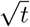 of diffusion (Appendix C). We write *σ*_*i*_ = *σ*(*t*_*i*_) for the width at the *i*th diffusion time, and *σ*_1_ for the finest. The width is a property of the operator, not a measured biological resolution. The lazy factor rescales the finest band’s amplitude, but not its correlation; only the estimated width moves (Appendix G). Equations (2) and (3) are the ruler the rest of the paper uses: **W** fixes how a field is smoothed, and *σ* converts each smoothing level into micrometres.

### 2.2 Domain oracle

#### Definition 1

(Partition-measurable predictor). *Let P* = {*P*_1_, …, *P*_*K*_} *partition the N spots. A predictor* **F** ∈ ℝ ^*N ×G*^ *is P-measurable if it is constant on every cluster of* P. *Equivalently, every column of* **F** *lies in* span 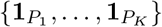; *we write* 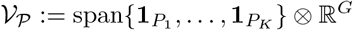 *for the set of such matrices*.

#### Proposition 1

(The doI:main oracle). *Let* **Π**_*P*_ *be the orthogonal projector onto V*_*P*_, *so that* 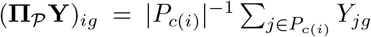 *with c*(*i*) *the cluster containing spot i. Then for every* P*-measurable* **F**,

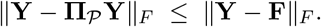

#### Proof

*V*_*P*_ is a linear subspace. The claim follows from the projection theorem in ℝ^*N ×G*^ under the Frobenius inner product.

The score of (1) is a correlation, not a squared error, so one more step is needed before the projection bounds the score.

#### Corollary 1

(Th e bound in the score). *For every* P*-measurable* **F** *and every gene* 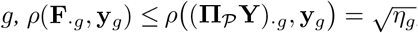, *where η*_*g*_ *is the share of the variance of* **y**_*g*_ *that lies between the clusters of P. Hence* PCC(**F, Y**) ≤ PCC(**Π**_*P*_ **Y, Y**), *and refining P cannot lower the right-hand side, since refinement enlarges V*_*P*_.

#### Proof.

*ρ* is invariant to **F**_*·g*_ *↦ a***F** _*·g*_ + *b***1** with *a >* 0 and **1** ∈ *V*_*P*_, so Cauchy–Schwarz in the centred subspace gives the bound, which the cluster mean attains (Appendix B).

**Π** _*P*_ **Y** is therefore the exact upper bound on the score for that partition: no predictor that sees only which cluster a spot lies in can beat it. Built from the measured **Y**, it is an oracle, and by construction it has no structure inside a cluster. A high oracle score therefore means that cluster identity alone suffices for a high score.

We take *P* from the image alone, by *K*-means on the principal components of a frozen pathology encoder. The clusters are spatially fragmented (Appendix K.4). The *oracle’s ratio* is its score as a fraction of a trained model’s, reported in per cent; each table states how the fraction is aggregated.

The oracle uses the measured expression of the sample it scores. To ask how much of that domain-level signal is learnable, a second predictor keeps the same image-derived partition and takes each cluster’s profile from training data instead. On the benchmark the other samples of the cohort are projected onto the scored sample’s feature basis and assigned to its clusters, and a cluster’s profile is the mean expression of the training spots assigned to it; on Xenium it is the mean over the training blocks. Every spot of the scored sample then receives its cluster’s profile, so the predictor is still constant within each cluster and never sees test expression. A variant that also learns the partition from the training data is reported in Appendix K.4.

Figure 1 previews the scalar and finest-band scores on two 900 *µ*m windows of the Visium HD colon sections, with the domain oracle at *K* = 20 and a ridge model on frozen features scored on each section’s held-out half over a 50-gene panel. The oracle is posterised into domains and the model is continuous, yet their correlations with the measurement differ by only 7.2% and 6.5% (0.827 against 0.892, 0.902 against 0.964). In the finest band of Section 2.4 they differ by factors of 5.4 and 3.6 (0.128 against 0.694, 0.200 against 0.714): the measurement stays textured everywhere, the oracle keeps only domain boundaries, and the model keeps little amplitude inside domains but places it correctly. That band holds 15.6% and 16.0% of the measured variance against 1.9–4.0% of either prediction. Section 3.3 tests the contrast on all 22 Xenium regions.

**Figure 1.**
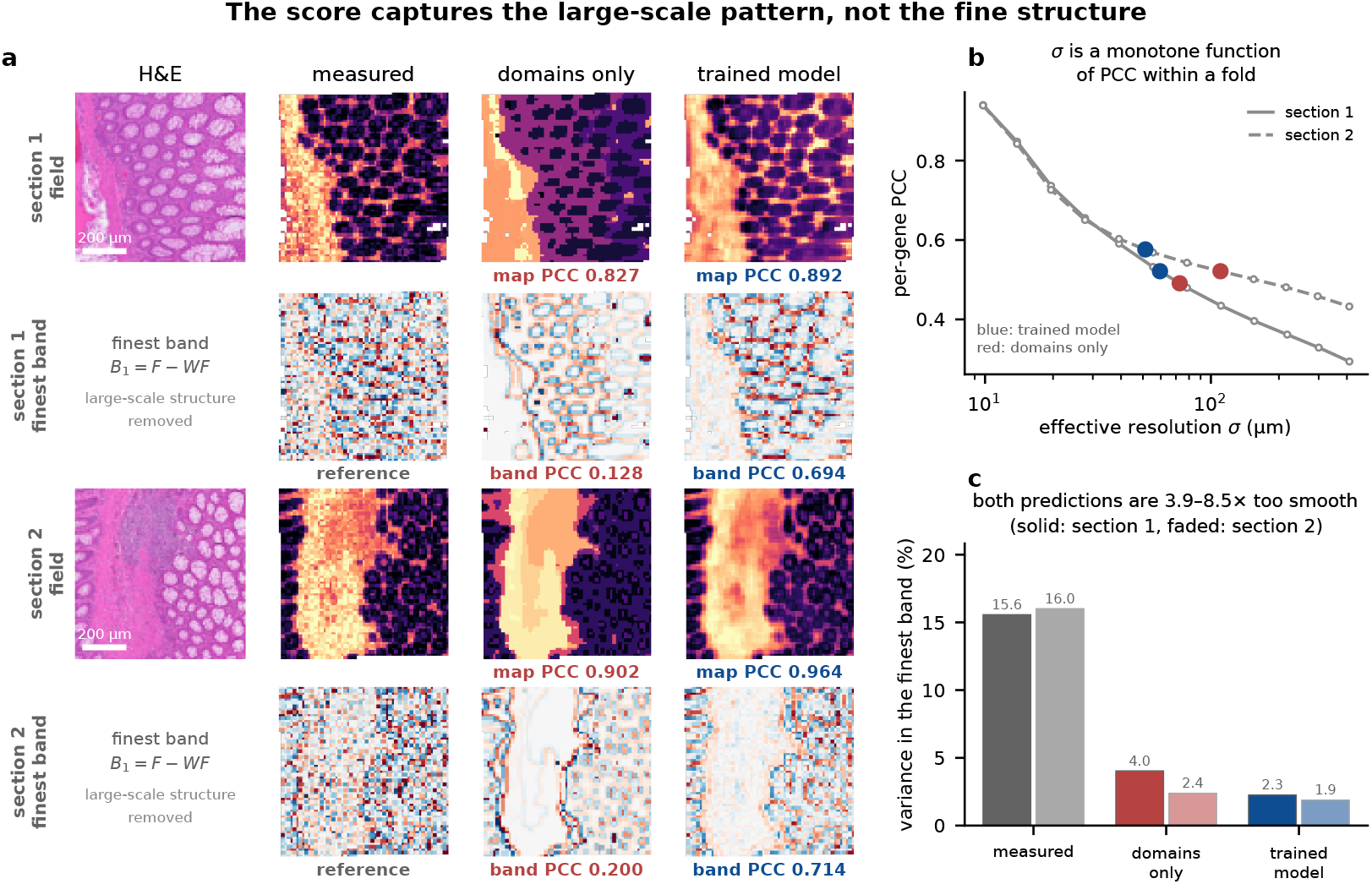
The window of each Visium HD colon section with the largest finest-band gap, among 900 *µ*m windows with at least 85% coverage and at least three image clusters where both predictions correlate at least 0.80 with the measurement and within 8% of the larger, enumerated on a 160 *µ*m step (13 of 433, 147 of 425). **(a)** For each window: H&E, the measured field and the two predictions (first row), and the same three maps with large-scale structure removed (second row). Maps are the panel’s first principal component; domains only averages the measured field within 20 image-derived domains; trained model is ridge on frozen features. The number under a map is its correlation with the measured map in the window. **(b)** The calibration curve of Eq. (4) for each section, with both predictors placed on it. **(c)** Share of variance in the finest band, per section.

### 2.3 Effective resolution

One natural way to give (1) units is to blur the truth until it scores what the model scores. Define the calibration curve

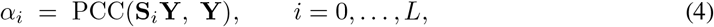

with **S**_*i*_ and *σ*_*i*_ from Section 2.1. It starts at *α*_0_ = 1. For a degree-weighted correlation we prove its monotonicity (Appendix B). For the Euclidean correlation we use, we check it instead: on all 22 Xenium regions and both Visium HD sections *α* is strictly decreasing at every one of the *L* = 12 steps, with a smallest step of 0.014. Let *α*( · ) be its interpolation in log *σ* over [*σ*_1_, *σ*_*L*_]; a score outside [*α*_*L*_, *α*_1_] is flagged rather than assigned a width. The effective resolution of a prediction is

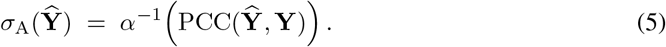

#### Proposition 2

(Degeneracy). *Fix the spots, the operator* **W**, *the diffusion schedule, the width estimates, the gene set and the target* **Y**. *Suppose α is strictly decreasing on the range of interest. Then σ*_A_ *is a strictly decreasing function of* PCC(**Ŷ, Y**) *alone. For any two predictions* **Ŷ**_1_, **Ŷ**_2_ *on the same fixed data*,

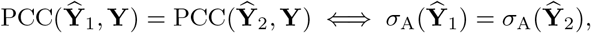

*and σ*_A_ *carries exactly the information in* PCC.

*Proof*. By (5) we have *σ*_A_ = *α*^*−*1^ ∘ PCC. Here *α*^*−*1^ depends only on quantities held fixed. A strictly decreasing *α* has a strictly decreasing inverse. Composing with a strictly decreasing bijection reverses the induced ordering and preserves the generated sigma-algebra.

The consequence is concrete: comparing two methods on the same data by effective resolution restates their correlation gap, and a figure that shows an intervention lowering the score and coarsening the resolution shows one fact twice. Across folds *α* changes with the tissue, so the same score can mean widely different physical scales (Appendix B).

### 2.4 Octave

Proposition 2 closes the obvious route: on fixed data, any calibration of the score into micrometres is a monotone reparameterisation of it. Two requirements therefore fix the construction below: it must not be a function of the score, and each band must carry a width in micrometres. The band scores are computed on filtered fields; the name Octave follows the dyadic diffusion times of Section 2.1, each band beyond the first spanning one octave.

#### 2.4.1 Band decomposition

Define the bands

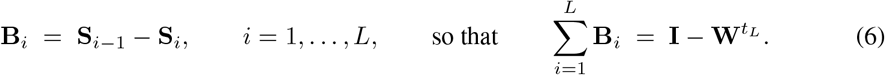

Band *i* keeps what survives *t*_*i−*1_ diffusion steps but not *t*_*i*_, and its width is *σ*_*i*_. The bands telescope to the identity minus the coarsest low-pass. Proposition 3 proves this.

##### Band-resolved score

The natural scale-resolved analogue of (1) is

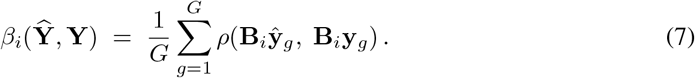

*β*_*i*_ is not a decomposition of PCC: the correlation is normalised inside each band, so no additive relation with fixed weights links {*β*_*i*_} to the scalar. We report each band score on its own.

#### 2.4.2 Estimand

Let **Ŷ**^mod^ be a trained model and **Ŷ**^dom^ = **Π**_*P*_ **Y** the domain oracle of Proposition 1. We write PCC(**Ŷ**) for PCC(**Ŷ, Y**) when **Y** is fixed. Define the *scalar shortfall* and the *fine-band shortfall* of the domain oracle relative to the model,

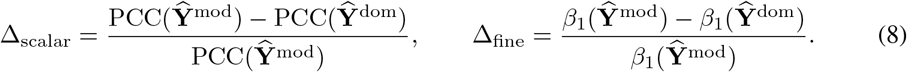

Both shortfalls are dimensionless, computed per region, and divided by the trained model’s value, which is bounded away from zero on every region we use (Appendix B). Our quantity of interest is the ratio Δ_fine_*/*Δ_scalar_: how much larger the normalised shortfall is in the finest band than in the score. The ratio compares two gaps. It is not a correction factor on the score. Its denominator can be small, so we report it as a per-specimen median over regions and then a median across specimens. The paired sign test of Δ_fine_ *>* Δ_scalar_ carries the inference.

## 3 Results

We first ask how much of the score coarse domain identity alone explains, on the public benchmark with the oracle of Proposition 1 and its learnable counterpart; then whether scoring each scale separately reveals what the scalar hides, with Octave on Xenium; and finally whether the findings hold across 57 encoders and what the benchmark’s own resolution can evaluate.

### 3.1 Data and protocol

#### Data

We use three sources. Twenty-two Xenium (Janesick et al., 2023) regions from 13 specimens, binned to 16 *µ*m: four breast specimens with three regions each, one lung specimen with two regions, and eight others with one. Two Visium HD (Oliveira et al., 2025) colon sections at 16 *µ*m, with four within-slide folds. And the HEST-Benchmark (Jaume et al., 2024): 72 samples, 10 cohorts, two platforms at a ∼100 *µ*m pitch, scored under its own protocol with 57 frozen encoders. They span 6M (Cai et al., 2026) to 1.1B parameters (Scalbert et al., 2026; Xu et al., 2024; Karasikov et al., 2025): forty-eight pathology encoders, eight of them distilled (Cai et al., 2026; Kaspar et al., 2026; Grashei et al., 2026; Filiot et al., 2025), eight general-purpose backbones (Oquab et al., 2024; Siméoni et al., 2026; Radford et al., 2021; Tschannen et al., 2025) and one biomedical vision– language model (Zhang et al., 2025). Inference runs at the specimen level for Xenium and the cohort level for the benchmark, with exact two-sided sign tests (Appendix D).

### 3.2 Coarse domain identity explains most of the score on the benchmark

Every encoder is scored on the same 72 samples under one protocol: we build the partition from its own features on the test sample and compare with a ridge model trained on the cohort’s other samples, with the penalty chosen on the other nine cohorts (the benchmark’s own setting is far too small and would flatter the oracle).

The learnable predictor of Section 2.2, whose cluster profiles come only from training data, reaches a median 82% of the trained model’s score (range 61–92% over encoders; Table 2), and learning the partition itself from training data as well leaves this at 79% (Appendix K.4).

The domain oracle bounds what that identity can earn. Table 1 gives all 57 encoders; Appendix I adds the paired differences and a summary by encoder size. In the cohort-level paired comparison the oracle scores above the trained model for 56 of 57 encoders, and the sign test is significant in the same 56; over each encoder’s sample mean the median lead is 47%. The 57 tests share ten cohorts, so we read the count as a consistency check; pooled over encoders, the median lead is positive in 10 of 10 cohorts (*P* = 0.0020). The one exception, OpenMidnight (Kaplan et al., 2025), is a single dominant feature coordinate; standardising the features removes it (Appendix J). The oracle is a bound: it is handed the measurements of the sample it scores. Refining the partition raises the bound, to a median lead of 62% at *K* = 50 and 90% at *K* = 200; as *K* → *N* the bound tends to one, so we fix *K* = 20, and Table 6 shows what that choice costs on Xenium. Corollary 1 guarantees the ordering only for nested refinements, so it is observed, not forced. Encoder size narrows the gap without closing it (Spearman *r*_*s*_ = −0.30 against parameter count over encoders, *P* = 0.024, and −0.37 over the 32 encoder families). The eight distilled encoders, at 6M to 87M parameters, sit with their 0.6B to 1.1B teachers rather than with same-size models trained from scratch; without them *r*_*s*_ = −0.60 over 49 encoders (*P <* 10^*−*4^) and −0.58 over 29 families (Appendix I). None of the eight end-to-end methods we ran under the benchmark protocol exceeds the bound. Six score below every one of the 57 encoders paired with ridge and win no cohort; DeepSpot’s four configurations sit inside the encoder range and tie its top only at their per-sample best, and TRIPLEX is scored per fold (Appendix H).

**Table 1:**
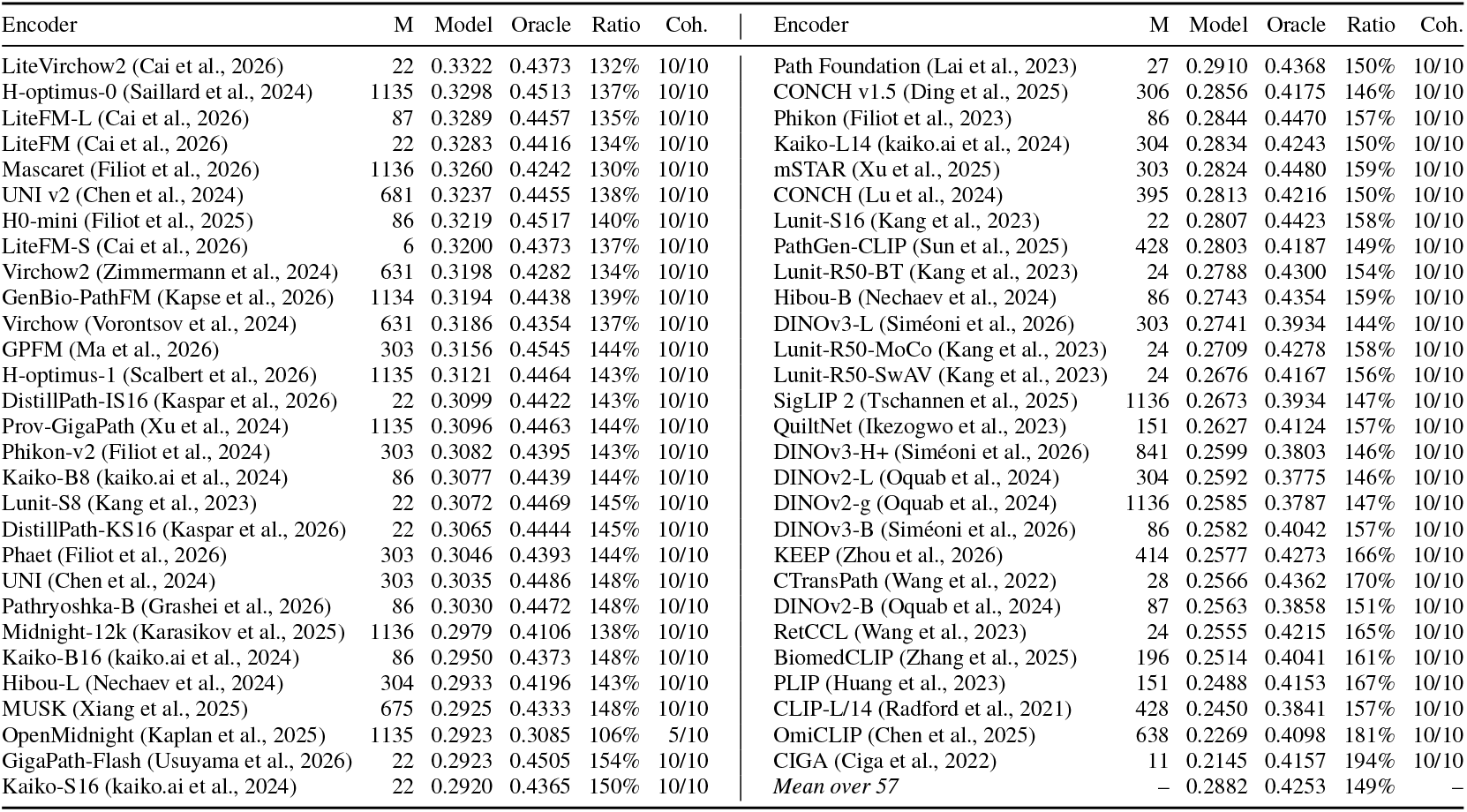
The domain oracle against a trained model on the public benchmark (72 samples, 10 cohorts, *K* = 20), for all 57 frozen encoders, ordered by the model’s own score. M is parameters in millions; Model and Oracle are mean PCC over samples; Ratio is Oracle over Model; Coh. counts cohorts in which the oracle leads; the last row means Model, Oracle and Ratio over the 57, so its Ratio is the mean of the per-encoder ratios. Appendix I adds paired differences and a summary by size

Whether the image is needed depends on the pitch. A partition of the spot coordinates knows nothing about the tissue; with measured means it scores 123% of the model on the benchmark, above it in a median 6 of 10 cohorts, but only 33% on Xenium at 16 *µ*m, where the learnable image partition keeps 79%. A random partition of the same sizes scores 0.08 and 0.01, the finite-sample optimism of measured means. Splitting the model’s own prediction on the image partition, with no test label, shows where its score lies: on Xenium its between-cluster part carries 79% of its PCC but a finest-band correlation of only 0.04 against 0.24, so what the finest band credits to the model is within-cluster structure (Appendix K.4). At ∼100 *µ*m the score rewards coarse spatial structure whatever its source; at 16 *µ*m the image is needed and the model wins.

### 3.3 Octave separates oracle and model as the scale gets finer

The protocol was fixed in advance: within-section spatial block cross-validation on a 16 × 16 grid of blocks, the 200 genes of highest variance (the benchmark’s own selection rule), and ridge on the frozen features of Hibou-L (Nechaev et al., 2024), with *P* a 20-cluster image partition. Under the scalar the two predictors look close: a median PCC of 0.428 for the oracle against 0.559 for the model, and a median scalar shortfall, formed per region, of 21% (Table 3). Band by band the picture changes: at the first band above the bin pitch, median width 19.8 *µ*m, the oracle’s shortfall is 2.37 times its scalar shortfall; at the grid limit, 10.3 *µ*m, the finest band reports 81% against the scalar’s 21%, a ratio of 4.04. Both ratios exceed one in all thirteen specimens (sign test *P* = 0.0002). At the four finest bands, of median width 10.3, 14.2, 19.8 and 27.4 *µ*m, the ratio is 4.04, 3.30, 2.37 and 1.45; the first three hold in 13*/*13 specimens and the fourth in 11*/*13 (*P* = 0.022). The finest band sits below the 16 *µ*m bin pitch, so we report the 20 *µ*m band first.

The profile (Figure 2a) levels off at about 11% beyond 100 *µ*m. The scalar’s 21% coincides with the band shortfall at *σ* ≈38 *µ*m. The reported score behaves like a measurement made at a few tens of micrometres, far coarser than the assay beneath it.

**Figure 2.**
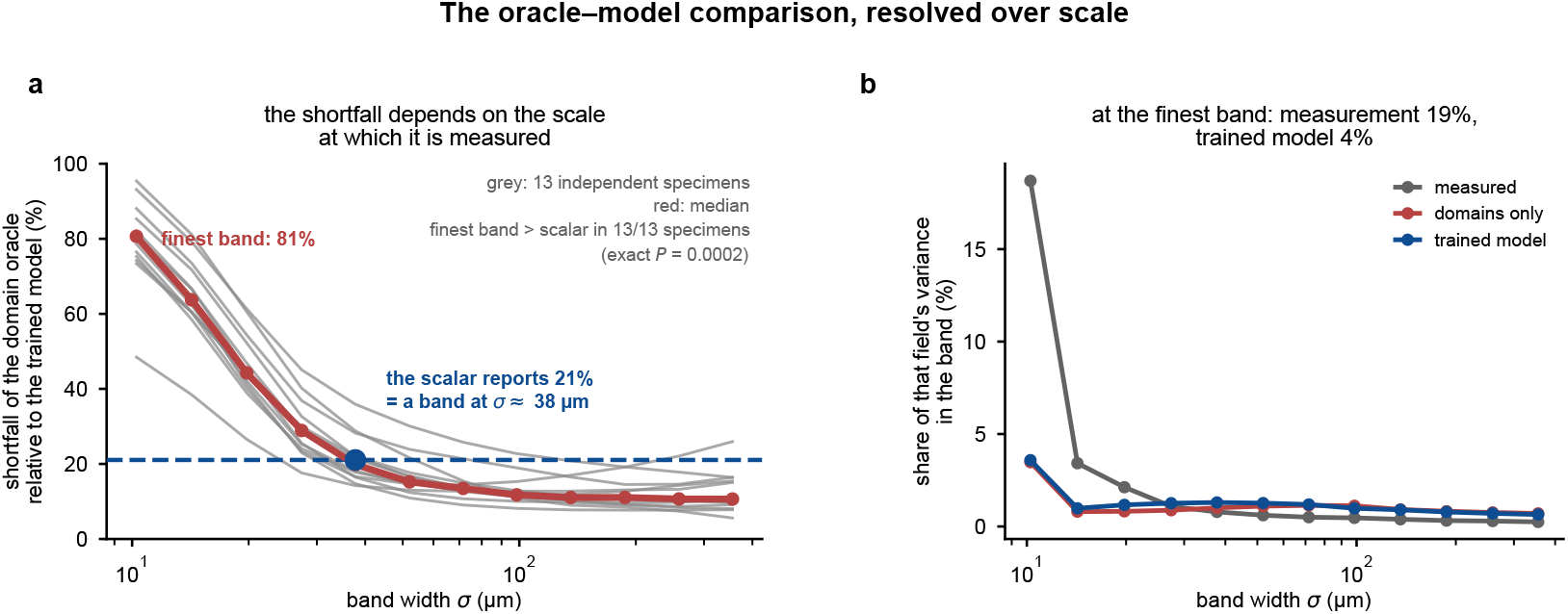
Twenty-two Xenium regions from thirteen specimens, binned to 16 *µ*m. **(a)** The shortfall of Eq. (8) against band width *σ*_*i*_. Grey lines are specimens, red their median, dashed the scalar; the specimen-level sign test is stated on the panel. **(b)** The share of each field’s own variance in each band

The pattern survives without access to test expression. The learnable predictor, whose cluster means come from the training blocks, matches the oracle’s finest-band correlation of 0.04 against the model’s 0.24, with an 81% shortfall and a ratio of 3.96, above one in 13 of 13 specimens; a cross-fitted oracle, with cluster means from one binomial half of the counts scored against the other, gives 3.90 (Appendix K.4). Across the thirteen specimens the finest band carries a median 19% of the measured field’s variance, 15% once the non-reproducible part is removed, against 1.5–6.9% for either prediction: both are too smooth by a median factor of 4.3 (Appendix K.3).

### 3.4 The finest band ranks fifty-seven encoders differently from the scalar

Across fifty-seven encoders under the same protocol (6M to 1.1B parameters; Appendix K.5), the scalar separates them little: scores run from 0.463 to 0.559, a range of 0.10. Finest-band correlations run from 0.0348 to 0.2381, a range of 0.20; best to worst that is a factor of 6.85 against 1.21. The ordering depends on scale: relative to the scalar ranking the finest band reorders 261 of the 1596 encoder pairs (Spearman 0.85), the *t* = 4 band 84 (0.98) and the *t* = 32 band 132 (0.96). A pair the scalar calls a tie is not one in the finest band: DINOv3-H+ and KEEP differ by 0.0039 in PCC and by 0.110 in *β*_1_, a 69% difference in finest-band fidelity. Resampling the 13 specimens (*B* = 2000) gives the finest-band ranking a Spearman correlation of 0.97 [0.92, 1.00] with the full-data ranking and never reverses DINOv3-H+’s lead over KEEP; 138 of the 261 reversed pairs keep their direction in at least 95% of resamples. The domain comparison generalises too: every encoder’s own oracle reaches 65–89% of its score, and the finest-band ratio stays between 2.4 and 5.9 for all fifty-seven. iStar (Zhang et al., 2024), an end-to-end super-resolution method run under a matched protocol, scores below ridge on both readouts in all eight specimens of the first data release (Appendix K.7).

### 3.5 The benchmark’s finest band is 50 *µ*m wide and cannot reach the gap

The benchmark samples at a ∼100 *µ*m pitch, and under the same construction its finest band has a median width of 50 *µ*m, half the spot pitch and coarser than the 10–30 *µ*m at which the oracle and the model separate on Xenium. Scored there, the oracle still leads, in a median 7 of 10 cohorts, by per-sample leads of 15% against 37% on the scalar (Appendix K.6). No score on this pitch can directly validate structure finer than that band, whatever the metric.

### 3.6 The direction survives every choice we varied

The direction holds at every ridge penalty, gene count and *K* of Appendix G and at each of 16 *K*-means seeds (Table 3, last row): the finest-band ratio exceeds one in 13*/*13 specimens throughout, ranging from 2.94 to 5.27, a span set by *K*. Default per-specimen ratios span 2.28 to 7.69. Two other operators agree: the symmetrically normalised walk gives a ratio of 3.70 and a Gaussian kernel 3.90 against the lazy walk’s 4.04, each above one in 13*/*13 specimens. The graph neighbourhood moves the finest width and the ratio (2.27 at *k* = 4, 3.67 at *k* = 12), and how genes enter the correlation moves it by at most 0.04 (Appendix G). The scoring geometry, by contrast, reverses a comparison: the same coarsening of a prediction is rewarded when the evaluation grid follows it and penalised when the grid stays fixed, in 15*/*15 regions and 7*/*7 specimens (*P* = 0.0156, Appendix F).

## 4 Related work

### Expression prediction from histology and its benchmarks

Many model families address this task (He et al., 2020; Pang et al., 2021; Zeng et al., 2022; Jia et al., 2024; Li et al., 2024; Xie et al., 2023; Wang et al., 2025b; Chung et al., 2024; Zhang et al., 2024; Wang et al., 2025a; Shi et al., 2024; Zhu et al., 2025), some built around multiscale image context (Li et al., 2026) or scored with spatial agreement (Pizurica et al., 2024), and some predicting below the spot pitch (Pang et al., 2021; Zhang et al., 2024; Shi et al., 2024). Public benchmarks standardise splits and report the mean per-gene correlation (Jaume et al., 2024; Mejia et al., 2024; Gindra et al., 2025); a seven-method assessment (Xie et al., 2026) adds domain detection. None of them asks what scale the correlation rewards.

### Benchmarks dominated by a non-target component

In single-cell perturbation prediction no deep model beat simple baselines (Ahlmann-Eltze et al., 2025), and the metric was inflated by the control population that defined the target (Nicol et al., 2026); the concern is older (Musgrave et al., 2020; Recht et al., 2019; Theis et al., 2016). Viñas Torné et al. (2026) answer it by scoring against the systematic variation a trivial predictor already captures; we resolve the correlation over scale, which says where a margin lives, not only whether it survives.

### Centre effects in frozen pathology encoders

Across twenty frozen encoders, 13.9%–55.4% of local embedding neighbourhoods are set by the medical centre, not by biology (Kömen et al., 2026), which one line of work answers by disentangling morphology from expression (Ostermaier et al., 2026). Those concern what the features encode, ours what the metric rewards.

### Multiscale analysis on graphs

Diffusion wavelets (Coifman & Maggioni, 2006) build on diffusion maps (Coifman & Lafon, 2006), and spectral graph wavelets (Hammond et al., 2011; Shuman et al., 2013) are close relatives; scale-space theory (Lindeberg, 1994) and MS-SSIM (Wang et al., 2003) do the same for images. We build such a filter on the graph of spots.

## 5 Discussion

A scale-blind score is reliable for the same reason it misses fine structure: it ranks methods mainly by coarse structure. The same gain can come from recognising domains or from recovering structure within them, and a scalar score cannot tell the two apart. The direction of the oracle–model comparison is stable; the level of the score is not. It moves substantially across Appendix G, and the grid changes of Appendix F reverse a comparison outright.

The same evaluation yields three quantities that complement the scalar: the learnable domain predictor’s fraction of the score, the finest-band correlation *β*_1_ and that band’s share of the measured variance. They show how much domain identity already explains, whether fine structure is placed correctly and how much of it the data contain (Table 2, Figure 1). Three copies of the measurement, degraded to the model’s PCC, score 0.088 (blurred), 0.179 (shifted) and 0.474 (noisy) in the finest band, against the model’s 0.238 (Table 16). Structure finer than a benchmark’s finest band can be validated only with data measured at that scale.

**Table 2:** Partition controls at *K* = 20: medians over 57 encoders of per-encoder sample medians (benchmark) and over 13 specimens of per-specimen medians (Xenium); “of model” is the median per-cohort or per-region ratio to ridge, formed inside each encoder or specimen. Table 10 uses each encoder’s sample means instead, putting the oracle at 147%. Full tables in Appendix K.4

| Predictor ($K = 20$) | Benchmark, $\sim 100 \mu\text{m}$ | | Xenium, $16 \mu\text{m}$ | |
| --- | --- | --- | --- | --- |
|  | PCC | of model | PCC | of model |
| Ridge on frozen features (trained model) | 0.268 | 100% | 0.559 | 100% |
| Image partition, measured means (domain oracle) | 0.414 | 140% | 0.428 | 79% |
| Image partition, means learned from training data | 0.246 | 82% | 0.427 | 79% |
| Image partition and means both learned from training data | 0.204 | 79% | 0.423 | 78% |
| Coordinate partition, measured means | 0.361 | 123% | 0.186 | 33% |
| Random matched-size partition, measured means | 0.080 | 21% | 0.011 | 2% |

**Table 3:** Shortfall of the domain oracle relative to the trained model, per specimen. Δ_fine_ *>* Δ_scalar_ in all thirteen (exact two-sided sign test *P* = 0.0002). Every column is a median over the specimen’s regions of its own per-region value, so a column is not the quotient of the printed values beside it. The last row repeats the whole computation at 16 *K*-means seeds (median ±s.d.; median and range for the ratio)

| Specimen | Regions | $\text{PCC}_{\text{mod}}$ | $\text{PCC}_{\text{dom}}$ | $\Delta_{\text{scalar}}$ | $\Delta_{\text{fine}}$ | Ratio |
| --- | --- | --- | --- | --- | --- | --- |
| Ovarian (Prime) | 1 | 0.672 | 0.600 | 10.7% | 82.2% | 7.69 |
| Skin | 1 | 0.616 | 0.531 | 13.7% | 95.4% | 6.95 |
| Cervical | 1 | 0.508 | 0.428 | 15.7% | 80.7% | 5.13 |
| Breast (Prime) | 1 | 0.505 | 0.424 | 16.1% | 81.4% | 5.06 |
| Lymph node | 1 | 0.534 | 0.421 | 21.1% | 85.3% | 4.05 |
| Lung | 2 | 0.586 | 0.477 | 18.9% | 73.4% | 4.04 |
| Breast S2 | 3 | 0.571 | 0.446 | 21.9% | 74.2% | 4.04 |
| Breast S1 | 3 | 0.596 | 0.474 | 19.6% | 78.6% | 3.90 |
| Breast S4 | 3 | 0.559 | 0.433 | 22.6% | 75.3% | 3.33 |
| Kidney | 1 | 0.423 | 0.292 | 31.1% | 93.1% | 2.99 |
| Breast S3 | 3 | 0.518 | 0.343 | 31.1% | 88.1% | 2.83 |
| Ovary (V1) | 1 | 0.569 | 0.394 | 30.7% | 76.5% | 2.49 |
| Prostate | 1 | 0.531 | 0.418 | 21.3% | 48.5% | 2.28 |
| Median |  |  |  | 21.1% | 80.7% | **4.04** |
| 16 seeds | | | | $21.0 \pm 0.1\%$ | $80.5 \pm 1.0\%$ | $3.93 [3.83, 4.05]$ |

### AI use statement

In this work we used a generative AI assistant for the following tasks: developing the conceptual framework, proposing and refining hypotheses and analyses, designing and giving feedback on experiments, formulating and drafting the statements and proofs of the propositions, implementing methods and data-processing code, and interpreting results. We did not use it to generate synthetic data. Translation, dataset cleaning beyond the released preprocessing code, and qualitative or thematic analysis are not applicable to this work. We also used it for creating and editing software and figure code, suggesting experimental parameters, drafting and editing the text, and locating candidate references.

We have reviewed all AI-assisted work. The authors checked the proofs line by line and withdrew what did not hold, and the choice of what to test and every decision about what the results license rested with the authors. We take responsibility for the final content of this work, including all text, claims and artifacts produced with the aid of generative AI.

### Reproducibility statement

The code is at https://github.com/QSong-github/OCTAVE. Table 4 lists the protocol settings, and Appendix E notes the one step that is not deterministic.

**Table 4:** Protocol settings. All values are fixed before any result is computed. Rows that name a platform apply to that platform (for Xenium, in Section 3.3 and Appendices G and K.4); the walk, the oracle partition and the finest band are shared by every platform. Appendix K.7 departs from these settings with a half split, stated there.

| Setting | Value |
| --- | --- |
| Spot grid (Xenium) | 16 $\mu\text{m}$ bins (the Visium HD bin size, about one cell across); $16 \times 16$ spatial blocks for cross-validation |
| Graph (Xenium) | symmetric 8-NN, edges $\leq 29 \mu\text{m}$ |
| Graph (benchmark, $\sim 100 \mu\text{m}$ pitch) | symmetric 6-NN, edges $\leq 1.15 \times$ median NN distance |
| Walk | $\mathbf{W} = (\mathbf{I} + \mathbf{D}^{-1}\mathbf{A})/2$ (lazy factor 1/2) |
| Diffusion times | $t_i = 2^{i-1}$; $t_L = 2048$, $L = 12$ (Xenium); $t_L = 1024$, 11 scales (benchmark) |
| Width estimator $\sigma(t)$ | 128 impulse sites (Xenium); 192 impulse sites (Visium HD); median over sites on Visium HD and the benchmark |
| Oracle partition | $K$-means, $K = 20$, on the first 50 PCs of the encoder’s test-section features; 4 restarts, seed 0 |
| Regression predictor | ridge on frozen features, training folds only; $\lambda = 10^4$ on Xenium and Visium HD, chosen on the other nine cohorts on the benchmark |
| Gene set (Xenium) | the 200 genes of highest variance, the benchmark’s own rule; variance taken over the whole region, as the benchmark does |
| Finest band | $\mathbf{B}_1 = \mathbf{I} - \mathbf{W}$, i.e. $t_0 = 0 \rightarrow t_1 = 1$ |

## Acknowledgements

Q.S. is supported by the National Institute of General Medical Sciences of the National Institutes of Health under award number R35GM151089. This work was also supported by NSF ACCESS “Discover” Allocation Award No. CIS261195 and No. CIS261339.

## A Limitations

Most of the *P* values we report are the minimum their design allows. Four of the thirteen Xenium specimens are breast sections from one release with unstated donor structure, so they may come from fewer than four donors. The thirteen span eight tissues, which fixes the direction of the effect but not its size elsewhere. The finest band has a reliability of *c* = 0.59, the lowest of the twelve bands (Appendix K.3), and the *K*-means partition behind the oracle reproduces only to the third decimal (Appendix E). The oracle’s level depends on the partition and rises with *K* in every setting we tried (Table 6).

## B Formal details

### B.1 Graph and operator

The occupied bins do not fill the grid, so degrees vary and a fixed stencil would not apply. We build the operator from the geometry. Let **A** be the adjacency of the *k*-nearest-neighbour graph on {*x*_*i*_} with *k* = 8, with an edge kept only when the two spots lie within 29 *µ*m of each other, and symmetrised as **A** ← max(**A, A**^*⊤*^). Let **D** = diag(max(**A1, 1**)), so that a spot with no neighbour within the radius has degree one, its row of **W** is 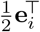 and its value decays on its own instead of mixing with neighbours. The radius is set from the 16 *µ*m binning, so that a spot reaches its eight neighbours on the grid and no further. The benchmark data lie on a coarser pitch and use their own graph, described in Appendix C.

The transition matrix **D**^*−*1^**A** is similar to the symmetric matrix **D**^*−*1*/*2^**AD**^*−*1*/*2^ through **D**^*−*1*/*2^. It is therefore diagonalisable with real eigenvalues. They lie in [ −1, 1], because the entries are non-negative and every row sums to at most one. A node with neighbours has row sum one. An isolated node has an empty row, so there 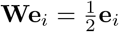 and **W1**≠ **1** (Appendix C gives the share of such nodes). Writing *ν* for an eigenvalue of **D**^*−*1^**A**, the lazy walk (2) has eigenvalues 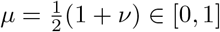.

Laziness is needed for the coarser bands. **D**^*−*1^**A** has eigenvalues near −1, and (**D**^*−*1^**A**)^*t*^ would alternate in sign as *t* grows, so a band **B**_*i*_ built from it would mix scales instead of separating them.

With every eigenvalue of **W** in [0, 1], **W**^*t*^ acts on each eigenvector by the non-increasing factor *µ*^*t*^. Each **S**_*i*_ is then a low-pass, and each **B**_*i*_ keeps the component that survives *t*_*i−*1_ steps but not *t*_*i*_.

### B.2 EXactness of the band decomposition

#### Proposition 3

(Telescoping). *With* 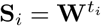, *t*_0_ = 0 *and* **B**_*i*_ = **S**_*i−*1_ − **S**_*i*_,

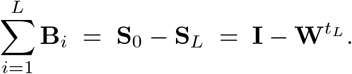

#### Proof.

The sum telescopes, and **S**_0_ = **W**^0^ = **I**.

The identity is exact for any schedule of diffusion times. It holds at the level of operators, so it holds for every field the operators act on. Two consequences matter for how the bands may be read.

First, the residual 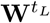 is the coarsest low-pass, and it carries whatever structure survives *t*_*L*_ steps. Our calibration curves run to *t*_*L*_ = 2048 over 12 dyadic scales. There *σ*_*L*_ has median 355 *µ*m and ranges from 265 to 418 *µ*m across regions. The residual is the field blurred to that width. It carries everything coarser, including the section mean.

Second, the **B**_*i*_ are not orthogonal projectors. They do not satisfy 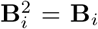, and **B**_*i*_**B**_*j*_ ≠ **0** for *i* ≠ *j*. They commute, because they are polynomials in **W**, but they do not form an orthogonal decomposition of the identity. This is why the band variances below do not sum to the total variance.

### B.3 Non-additivity of the band scores

#### Remark 1.

*There is no relation of the form* PCC = ∑_*i*_ *c*_*i*_*β*_*i*_ *with weights c*_*i*_ *that do not depend on the fields*.

The reason is visible in (7). Each *β*_*i*_ divides by 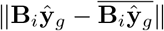 and by the matching norm of the measurement. The normaliser is computed inside band *i* and differs from band to band. A band that holds one per cent of the field’s variance and a band that holds half of it are both rescaled to unit norm before the correlation is taken. The band scores therefore lose the amplitudes that any additive identity would need.

We compare predictors one band at a time and never sum or average the {*β*_*i*_} into a version of PCC.

Proof of proposition 1

Fix the partition *P* = {*P*_1_, …, *P*_*K*_} and let *V*_*P*_ ⊂ ℝ^*N ×G*^ be the set of matrices constant on every cluster of P in every column. This set is closed under addition and scalar multiplication, so it is a linear subspace. Equip ℝ^*N ×G*^ with the Frobenius inner product ⟨**U, V**⟩_*F*_ = tr(**U**^*⊤*^**V**). The projection theorem in a finite-dimensional inner product space gives a unique minimiser of ∥**Y** − **F**∥_*F*_ over **F** ∈ *V*_*P*_, namely the orthogonal projection **Π**_*P*_ **Y**.

The projection has a closed form. The indicators 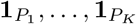 are orthogonal and span the subspace of vectors constant on every cluster, in each column, so

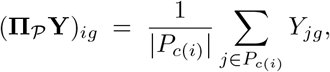

with *c*(*i*) the cluster containing spot *i*. The oracle is the cluster-wise mean of the measurement. It uses **Y**, which is why it is an oracle.

**Proof of Corollary 1**. Fix a gene and write **y** for its column, **f** for the column of a *P*-measurable **F**, and **u**_0_ = **u** − *ū***1** for centring. TI:he correlation *ρ*(**f**, **y**) = ⟨**f**_0_, **y**_0_⟩*/*(∥**f**_0_∥∥**y**_0_∥) is unchanged by **f** → *a***f** + *b***1** with *a >* 0. Because 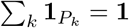, the constant vector lies in 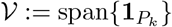, so **f**_0_ lies in *V* _0_ := *V* ∩ **1**^⊥^. Let Π_0_ be the orthogonal projector onto _0_. Then ⟨**f**_0_, **y**_0_⟩ = ⟨**f**_0_, Π_0_**y**_0_⟩ ≤ ∥**f**_0_∥ ∥Π_0_**y**_0_∥ by Cauchy–Schwarz, with equality, when Π_0_**y**_0_ ≠ **0**, if and only if **f**_0_ is a positive multiple of Π_0_**y**_0_. Hence *ρ*(**f**, **y**) ≤ ∥Π_0_**y**_0_∥ */* ∥**y**_0_∥ . The projector onto *V* fixes **1** and preserves the mean. So 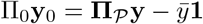, the centred cluster-mean predictor. The bound is therefore attained by **Π**_*P*_ **Y**. The quantity ∥Π_0_**y**_0_∥ ^2^*/* ∥**y**_0_∥ ^2^ is the share *η*_*g*_ of the variance of **y** that lies between clusters. Averaging over genes gives the statement for PCC. If *P*^*′*^ refines *P* then 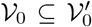 and 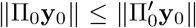, so the bound cannot fall. If all cluster means of a gene coincide, Π_0_**y**_0_ = **0**, every non-constant *P*-measurable predictor scores exactly 0, and the oracle’s own correlation is 0*/*0. We score such a gene as 0 on Xenium and Visium HD, and the benchmark pipeline omits it from the mean. No *P*-measurable predictor can exceed 0 on such a gene.

Two limits fix how the bound may be read. If *K* = 1 every gene has zero between-cluster variance and the score is 0 under that convention. If *K* = *N* every cluster is a single spot, **Π**_*P*_ = **I**, and the bound is vacuous. Between the two, the bound rises with *K* on the independent partitions of Table 6.

### B.5 Proof of proposition 2

Fix the spots, the operator **W**, the diffusion schedule, the width estimates, the gene set and the target **Y**. Every quantity entering the calibration curve (4) is then fixed, so *α* is a fixed function of the scale index, and its interpolation in log *σ* is a fixed function *α* : [*σ*_1_, *σ*_*L*_] → ℝ.

Assume *α* is strictly decreasing on the range of interest. A strictly decreasing continuous function on an interval is a bijection onto its image, and its inverse *α*^*−*1^ is strictly decreasing as well. By (5),

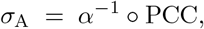

a composition of the scalar score with a fixed strictly decreasing bijection. Strict monotonicity gives the stated equivalence in both directions, and reverses the induced order. Because *α*^*−*1^ is a bijection, the *σ*-algebra generated by *σ*_A_ equals the one generated by PCC. No function of *σ*_A_ can separate two predictions that PCC does not separate.

We prove monotonicity for the degree-weighted correlation. With ⟨**u, v**⟩_**D**_ = **u**^*⊤*^**Dv** the operator isself-adjoint with spectrum in [0, 1] and, on a graph without isolated nodes, fixes **1**, so for **y**_0_ ⊥_D_ **1** with eigen-coefficients 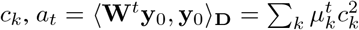 and the weighted correlation of **W**^*t*^**y**_0_ with **y**_0_ is 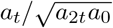 . Writing 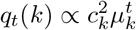 for the tilted weights, 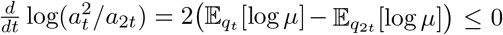, because *q*_2*t*_ places more weight on larger *µ* than *q*_*t*_ does. The score of (1) uses the Euclidean correlation instead. On an irregular graph it need not be exactly monotone. On all 22 Xenium regions and both Visium HD sections it is. Every one of the 12 steps of every curve decreases, and the smallest step is 0.014.

#### Non-degeneracy of *σ*_A_ across folds

The calibration curve *α* changes from fold to fold, because different tissue has different spatial autocorrelation. Across folds *σ*_A_ is therefore not a monotone image of PCC. The same score can then mean widely different physical scales. One pair from an earlier experiment makes this concrete. A model transferred across patients scored 0.6954 at *σ*_A_ = 74 *µ*m, and the same method transferred from tumour to normal colon scored 0.6897 at 183 *µ*m, within 0.006 in PCC of the first. That run used four Visium HD colon sections, two of them outside the three sources of Section 3.1, and is not part of the results reported here. The tumour-to-normal side rests on one test section.

### B.6 Estimand

Both shortfalls in (8) divide by the trained model’s score rather than by the oracle’s, which can be small. The model’s score is bounded away from zero on every region we use. Across the 22 Xenium regions its minimum is 0.423 in PCC and 0.124 in *β*_1_.

The ratio of the two shortfalls does divide by a difference, so we never average it over regions or specimens. We form the ratio per region, take the median within a specimen, and take the median across specimens. Aggregating in the other order, by taking the median of each shortfall within a specimen first and dividing after, changes the value. On our data the two orders give 4.04 and 3.88, respectively. We use the per-region order throughout, including in Table 3.

## C Protocol details

### Multiscale operator

On the 16 *µ*m data the graph is a symmetric 8-nearest-neighbour graph on spot coordinates with edges truncated at 29 *µ*m. Bins with no neighbour within that radius are isolated. They are a median 0.6% of bins per region and at most 3.0%. On an isolated bin **W** has diagonal 1*/*2 and no off-diagonal mass, so it keeps half that bin’s own value and a band does not cancel a constant there, which is harmless because predictions carry no free offset. We keep these bins. Under the default cut only the four orthogonal neighbours at 16 *µ*m and the four diagonal ones at 22.6 *µ*m survive. The *k* = 12, 40 *µ*m graph of Appendix G reaches the next ring at 32 *µ*m and connects most of the isolated bins. It gives the same conclusion. The diffusion times are powers of two up to 2048. We evaluate (3) with 128 impulse sites drawn without replacement. Over three orders of magnitude in *t, σ* grows as *t*^0.43^ to *t*^0.49^ across the Xenium regions. Widths at adjacent diffusion times differ by a factor of 1.30 to 1.41. That is close to the 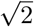 of exact diffusion. We therefore interpolate in log *σ*.

### Platform differences

Three details are stated here so that the numbers can be reproduced. First, the benchmark samples lie on a ∼100 *µ*m pitch, and their graph is a symmetric 6-nearest-neighbour graph with edges truncated at 1.15 times the median nearest-neighbour distance; diffusion times run up to 1024 over 11 scales. Second, on the Xenium data the width of (3) is estimated as written. It is the mean over impulse sites of the second moment about the site. On the Visium HD and benchmark data it is the median over impulse sites of the variance about the kernel’s centroid. Visium HD uses 192 sites. On the sixteen Xenium regions of the first data release, where both estimators were run, the two estimators agree to within 3% at the median over regions and scales, and to within 12% in the worst region at the coarsest scale. Third, the inverse of the calibration curve is taken in log *σ* on all three sources. The only quantity these three choices touch that we compare across platforms is the band width, where the benchmark’s 50 *µ*m is a factor of 1.7 to 5 above the 10–30 *µ*m at issue on Xenium. None of these choices moves *σ* by anything of that size. A further difference is intrinsic to the measurement. Two of our sources are sequencing-based: Visium HD and the Visium samples of the benchmark. On them, lateral diffusion of captured mRNA makes the effective resolution of the measurement itself coarser than the spot pitch. The amount varies across platforms and tissues (You et al., 2024). Xenium detects transcripts in situ and is not subject to this effect. This is one reason our finest-scale claims rest on the Xenium regions. The effective resolution of Section 2.3 is a property of a predictor relative to the measurement, not of the platform.

### Band homogeneity

Pooling a band across samples is only meaningful if the band means the same physical scale in each. The finest band does. Across all 72 benchmark samples *σ* spans 50.0–50.2 *µ*m, a ratio of 1.00. The two coarsest bands do not. There *σ* spans 641–1109 *µ*m and 758–1488 *µ*m, ratios of 1.7 and 2.0. We exclude those bands from every pooled benchmark statistic. On the Xenium regions we test at the finest band, where *σ* spans 9.9–10.6 *µ*m, a ratio of 1.07. The per-specimen profile of Figure 2a uses all twelve bands, the coarsest of which spans 265–418 *µ*m.

### Predictors

The trained model is ridge regression on frozen encoder features, at *λ* = 10^4^ on Xenium and Visium HD and, on the benchmark, at a penalty chosen on the other nine cohorts (Section 3.2). We fit it on training folds only. The domain oracle runs *K*-means on the first 50 principal components of the same features. It then replaces each spot by the measured mean of its cluster. The two predictors share the same folds and gene set.

### Consolidated settings

Table 4 collects the protocol choices that every result in this paper holds fixed. Appendix G varies the ridge penalty, gene count, *K*, graph and walk one at a time and reports how the headline quantities move.

### Reading the bands

A diffusion band is not a Fourier band. **W**^*t*^ averages a field over a neighbourhood that grows with *t*. So **S**_*i*_*f* is the field seen at width *σ*(*t*_*i*_). The band **B**_*i*_*f* = **S**_*i −* 1_*f −* **S**_*i*_*f* is the part of *f* still visible at width *σ*(*t*_*i*_−_1_) that width *σ*(*t*_*i*_) has already averaged away. The finest band therefore holds the spot-to-spot contrast that one diffusion step already removes, and the residual holds everything coarser than *σ*(*t*_*L*_). The bands and the coarsest low-pass sum to the identity (Proposition 3), but the bands are not orthogonal, since **W** is not a projection, and their variances do not add up to the total variance. So for each band we report the correlation of two band-filtered fields, not a share of explained variance. And *β*_*i*_ is the fidelity at width *σ*_*i*_. The dyadic schedule makes successive widths grow by a roughly constant factor, the 1.30 to 1.41 reported above. That is what lets the sequence of *β*_*i*_ values be read as a profile over scale.

## D Statistical conventions

Every sign test runs at the level of independent specimens for Xenium, or cohorts for the public benchmark, never at the level of regions or samples. The reason is simple: of the 22 Xenium regions, 14 come from 5 specimens, and 47 of the 72 benchmark samples come from two cohorts. Counting regions or samples would be pseudoreplication in the sense of Hurlbert (1984). The correlations against parameter count take the encoder or the family as their unit. We use exact two-sided sign tests. The minimum attainable *P* value is 0.0156 at *n* = 7, 0.0078 at *n* = 8, 0.0020 at *n* = 10 and 0.0002 at *n* = 13.

The shortfalls of (8) and their ratio are formed and aggregated as in Appendix B. Every other relative quantity is formed per cohort or per region and then aggregated, so it is not the quotient of the levels we print.

## E Reproduction

Two details affect reproduction. Ridge regression is deterministic given the folds. Rerunning it returns identical numbers, and we checked this in the 16 regions of the first release, where stored scores exist to compare against. The *K*-means partition behind the domain oracle is not deterministic in the same way, because it depends on the BLAS thread count. On one Visium HD fold, partitions computed at 8 and 32 threads differ, with an adjusted Rand index of 0.83, and about 11% of spots change cluster; the resulting scores differ in the third decimal. We therefore pin the thread count and check the reproduced score against the stored one. Rerunning the oracle at the stored seed and thread count reproduces the stored score exactly in 7 of those 16 regions, and to within 4.6 × 10^*−*5^ in PCC in the other nine. That discrepancy comes from a neighbouring local optimum of the same partition problem. In all 16 it is smaller than the seed-to-seed spread of the same quantity. The across-seed standard deviation has median 1.7 × 10^*−*3^ in PCC. The seed spread, not the discrepancy, is the uncertainty of an oracle score.

## F Evaluation grid

We hold the predictions fixed and change only the geometry on which they are scored. The same coarsening of the prediction is then rewarded or penalised, depending on whether the evaluation grid follows the prediction or stays fixed. Table 5 gives both settings. The sign reverses in 15*/*15 regions and 7*/*7 specimens (*P* = 0.0156).

**Table 5:** One coarsening of the prediction, from 8 to 64 *µ*m, scored two ways. The predictions are identical. Only the grid on which they are scored changes. Fifteen Xenium regions from seven specimens, the set available when this analysis was run; the cervical region and the six regions added after it are not included. Percentages are changes in the score from the 8 *µ*m to the 64 *µ*m prediction, each row on its own grid: the grid that follows the prediction for the first row, the fixed 16 *µ*m grid for the second

| Evaluation grid | Median (%) | Range (%) | Regions > 0 | Specimens > 0 |
| --- | --- | --- | --- | --- |
| Follows the prediction | **+42.8** | +24.0 to +81.2 | 15/15 | 7/7 |
| Fixed at 16 $\mu\text{m}$ | **-17.2** | -28.9 to -10.0 | 0/15 | 0/7 |
| Difference (pp) | 60.0 pp |  | 15/15 reverse | 7/7 reverse |

The slope of *σ*_A_ against the score is also protocol-dependent. Under three protocols, +0.01 PCC corresponds to a −12.0%, −17.6% or −6.2% change in *σ*. The fit within each protocol is good, with *R*^2^ = 0.89–0.99. The rate spans a factor of 2.8 across protocols. It also reverses sign, to +2.0%, when the only intervention is coarsening the output grid.

Figure 3 plots Table 5 and adds panels that a table cannot carry.

**Figure 3.**
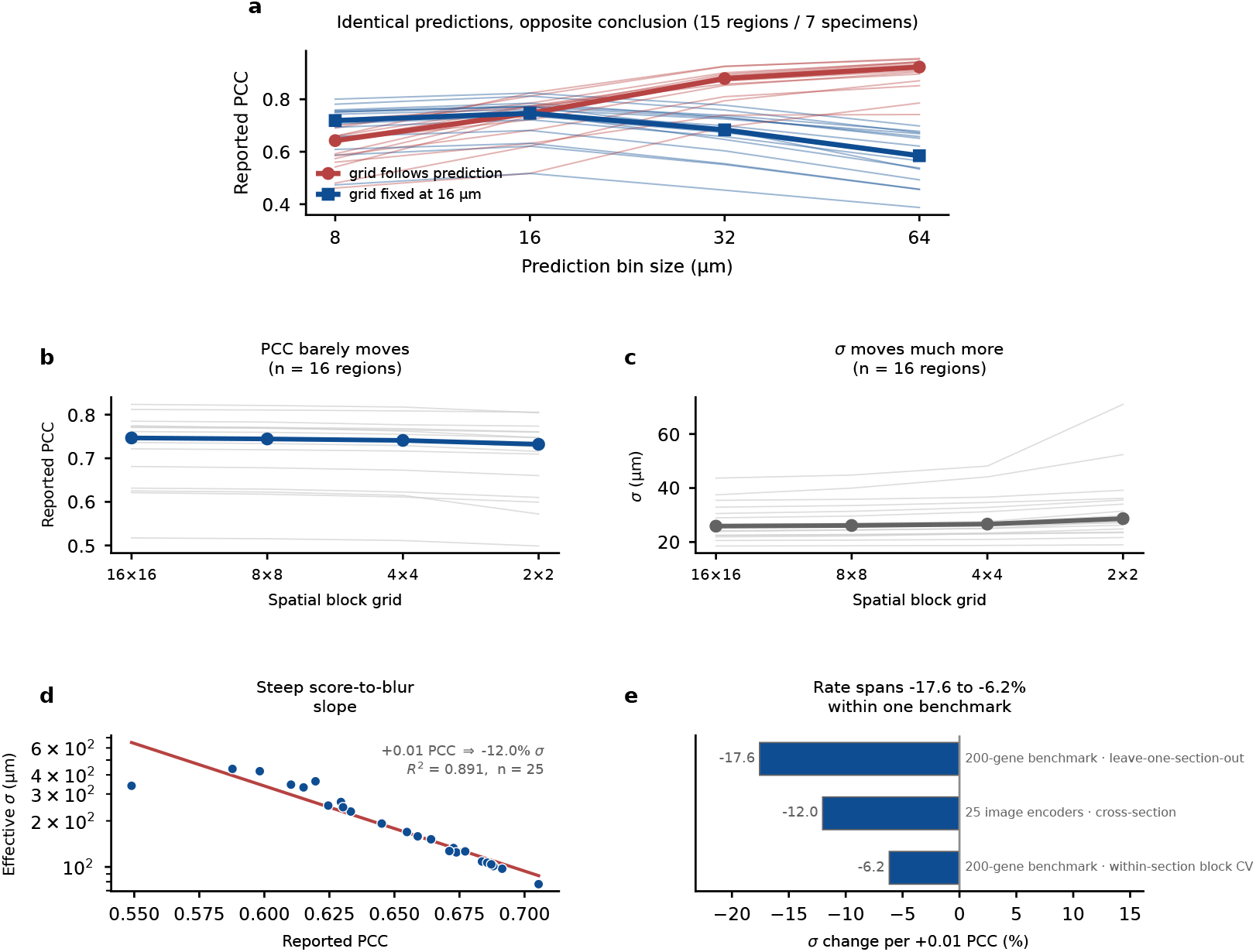
Table 5, plotted. Panel a uses the fifteen regions of Table 5; panels b and c the sixteen regions of the first release. **(a)** Reported PCC against prediction bin size under both evaluation grids. **(b–c)** The spatial block grid swept from 16 × 16 to 2 × 2, and the resulting change in PCC and, through each region’s calibration curve, in *σ*_A_. **(d–e)** The slope of *σ*_A_ against the score, by protocol; the encoder series pools the 25 encoders that were scored across sections on Visium HD, hence its *n* = 25

Under the fixed grid the reported score peaks at a prediction bin of 16 *µ*m (Figure 3a). Panels b and c sweep the spatial block cross-validation grid from 16 × 16 to 2 × 2 on the 16 regions of the first release, holding the rest of the protocol fixed. Panels b and c, and the two method series of panel e, come from an earlier 200-gene run that their labels call a 200-gene benchmark; it is not the public benchmark of Section 3.2, and its within-section scores sit above those of Section 3.3. The percentages below are formed within each run and are unaffected by that offset. Across the sweep, the reported score falls by 1.95%. Read through each region’s calibration curve, the same fall is a 10.70% coarsening of the effective resolution. The curve has a local slope of about 5.5 there. By Proposition 2 the 10.70% restates the 1.95% in the units of that curve. A choice rarely stated in papers moves the score. The move is small on the correlation scale and clear on the scale of micrometres.

## G Hyper-parameter sensitivity

Table 6 varies one choice at a time and holds the rest at their defaults, shown in bold. Each entry is the median over 13 specimens of the per-specimen median over regions, which is the aggregation used in the main text. Each entry is also the median over three *K*-means seeds. The gene axis stops at 200 because larger panels are not available in every region. The three default entries agree to the last digit reported, as they must, and at 3.90 they sit inside the sixteen-seed interval of Section 3.3, whose median is 3.93.

**Table 6:** One choice varied at a time. Oracle / model is the domain oracle’s score as a percentage of the trained model’s. Specimens counts those with Δ_fine_ *>* Δ_scalar_; every row rests on the same thirteen specimens, so the sign test gives 0.0002 wherever that count is 13/13. Defaults in bold

| Axis | Setting | Oracle / model | $\Delta_{\text{scalar}}$ | $\Delta_{\text{fine}}$ | Ratio | Specimens | $P$ |
| --- | --- | --- | --- | --- | --- | --- | --- |
| Ridge $\lambda$ | $10^2$ | 78.6% | 21.4% | 81.1% | 3.87 | 13/13 | 0.0002 |
| | $10^3$ | 78.5% | 21.5% | 81.1% | 3.85 | 13/13 | 0.0002 |
|  | **$10^4$** | **78.9%** | **21.1%** | **80.7%** | **3.90** | **13/13** | **0.0002** |
| | $10^5$ | 81.1% | 18.9% | 77.8% | 4.21 | 13/13 | 0.0002 |
| | $10^6$ | 86.2% | 13.8% | 71.9% | 5.14 | 13/13 | 0.0002 |
| Genes | 50 | 83.2% | 16.8% | 79.8% | 4.48 | 13/13 | 0.0002 |
|  | 100 | 81.8% | 18.2% | 80.1% | 4.18 | 13/13 | 0.0002 |
|  | **200** | **78.9%** | **21.1%** | **80.7%** | **3.90** | **13/13** | **0.0002** |
| Clusters $K$ | 5 | 69.2% | 30.8% | 87.6% | 2.94 | 13/13 | 0.0002 |
|  | 10 | 75.4% | 24.6% | 86.4% | 3.39 | 13/13 | 0.0002 |
|  | **20** | **78.9%** | **21.1%** | **80.7%** | **3.90** | **13/13** | **0.0002** |
|  | 50 | 82.7% | 17.3% | 77.3% | 4.50 | 13/13 | 0.0002 |
|  | 100 | 85.4% | 14.6% | 70.5% | 4.94 | 13/13 | 0.0002 |
|  | 200 | 87.7% | 12.3% | 65.7% | 5.27 | 13/13 | 0.0002 |

### Graph, walk and gene handling

Table 7 varies the graph that defines the walk and the way genes enter the finest-band correlation. Shrinking the neighbourhood to the four axial neighbours (*k* = 4, cut 17 *µ*m) reduces the finest-band shortfall from 81% to 48%. The one-step band then holds only the axial contrast, and its width falls to 7.1 *µ*m. The ratio drops to 2.27. It stays above one in 13*/*13 specimens. Enlarging the neighbourhood to *k* = 12 (cut 40 *µ*m) gives a ratio of 3.67. The lazy factor leaves the finest band’s correlation exactly unchanged. For any lazy factor *a* ∈ (0, 1), **B**_1_ = **I** − **W** = (1 − *a*)(**I** − **D**^*−*1^**A**). Correlation ignores that rescaling. Only the estimated width moves, from 10.3 to 7.3 *µ*m. Coarser bands do change with *a*. The symmetrically normalised walk and a Gaussian kernel give ratios of 3.70 and 3.90, each above one in 13*/*13 specimens (Appendix K.4). Dropping the quarter of genes with the smallest finest-band variance share moves the ratio by at most 0.04. Weighting genes by that share does the same. A 5% floor on that share removes at most 2 genes in any region.

**Table 7:**
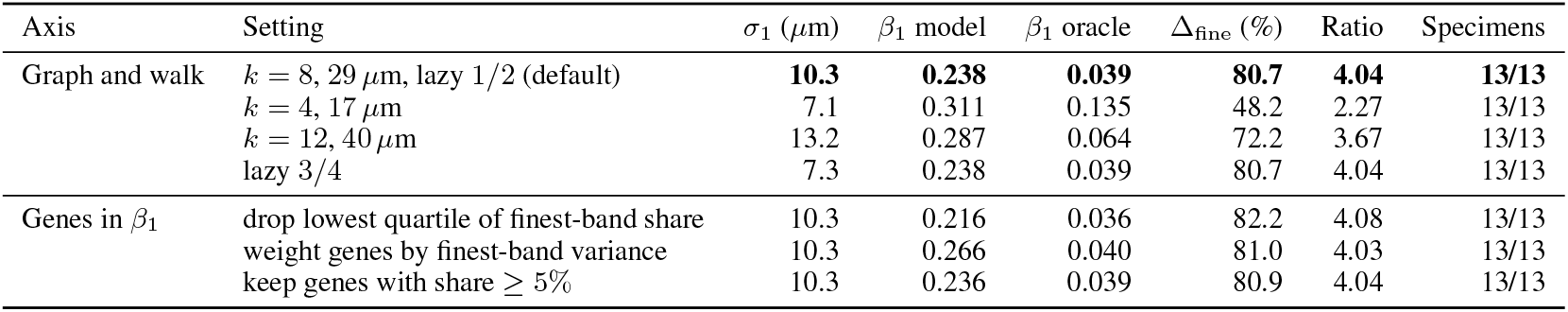
Operator and gene-handling sensitivity of the finest-band result, one choice varied at a time. *σ*_1_ is the median finest-band width over regions; the other columns follow Table 6, except that these entries come from a single *K*-means seed rather than the median over three. Δ_fine_ is formed per region before the medians, so it is not the normalised difference of the two *β*_1_ columns beside it. Ratio is Δ_fine_*/*Δ_scalar_ formed per region, then reduced to its median within each specimen and then across specimens, so it is not the quotient of the columns shown; Specimens counts those with ratio above one.

## H Eight end-to-end methods against frozen encoders with ridge

Our trained model throughout is ridge regression on frozen encoder features. To test whether a model trained end to end for this task scores higher, even above the oracle, we ran eight such methods (Pang et al., 2021; Zeng et al., 2022; Xie et al., 2023; Wang et al., 2025b; Li et al., 2024; Jia et al., 2024; Nonchev et al., 2025; Chung et al., 2024) on the same benchmark, under the protocol of Section 3.2, with the benchmark’s own splits, its 50 genes and the same expression target. Each model is the authors’ own code with its published hyperparameters. We changed the data source and two lines of code, one that current libraries reject and one that dominates runtime, both recorded below. Hist2ST, HGGEP and THItoGene, whose losses need raw counts, received them alongside the log-transformed target, as their own code does. A subsampling step caps a slide at 4,000 spots. It touches 19 of the 72 slides: on 17 it removes at most a quarter of the spots, on the two largest IDC slides half and four fifths. It thins over a block grid laid on the spot array and, where a slide still exceeds the cap after thinning, truncates in block order, so the retained spots need not cover the whole footprint. The three graph methods, Hist2ST, HGGEP and THItoGene, build their adjacency after thinning, with the authors’ own rule of four grid neighbours; a spot left with no neighbour is joined to its nearest spot. The cap applies to the three graph methods, to HisToGene and to TRIPLEX; ridge, BLEEP, HECLIP and DeepSpot are scored on every spot.

Table 8 gives the result for the six earlier methods and for DeepSpot (Nonchev et al., 2025), which, like our own predictor, builds on frozen pathology features. The six all score below every one of the fifty-seven frozen encoders paired with ridge, and none beats the strongest encoder in any cohort. DeepSpot’s authors name no default configuration, so we ran it on two backbones, UNI and H-optimus-0, at two training lengths, the ten epochs of their notebook and the 500 steps their paper implies, and report each configuration as its own row. The four average 0.296 to 0.329, below the strongest encoder’s 0.332 and within a hundredth of ridge on the same two backbones, and each trails the strongest encoder in seven or eight of the ten cohorts. Even with each sample’s best of the four, 0.336, DeepSpot only ties the strongest encoder ( −0.002 ±0.007 across cohorts, 5 of 10 won). A frozen encoder with a linear head is therefore a strong baseline for this task, and a method built for sub-spot detail lands in the same place.

**Table 8:**
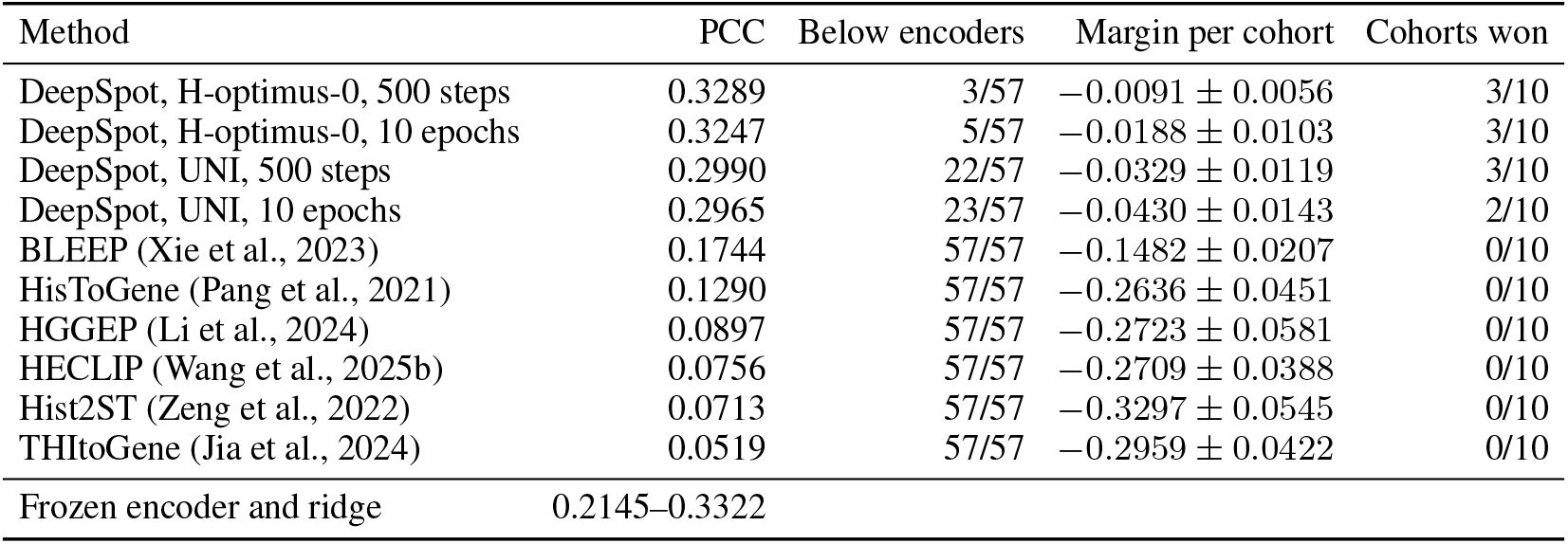
Seven end-to-end methods on the same 72 samples and 10 cohorts, against the fifty-seven frozen encoders paired with ridge. PCC is the mean over samples. The margin per cohort is against the strongest encoder, in PCC units, as the mean and its standard error over the ten per-cohort median paired differences. DeepSpot’s authors (Nonchev et al., 2025) fix no default configuration, so each of its four configurations, two backbones at two training lengths, is a row of its own

### A recent multi-resolution method

TRIPLEX (Chung et al., 2024) reads three views of each spot: the spot tile, a 1120-pixel neighbourhood split into 5 × 5, and attention over every spot in the section. It is the strongest end-to-end competitor we ran, and it needs its own comparison because its score is a per-fold mean rather than a per-sample one. We ran it with the authors’ code and published settings on two feature extractors: the self-supervised ResNet-18 of their own experiments, and UNI. Their evaluation selects the checkpoint by test-fold correlation, so we report their number under their own rule; our ridge penalty is chosen on the other nine cohorts and never sees the test fold. Table 9 gives the result. On the same features TRIPLEX is the better predictor: it beats ridge in 8 of 10 cohorts on the ResNet-18 features and in 6 of 10 on UNI, where its mean is 0.377 against ridge’s 0.372. One of those ten is a partial comparison, marked in the table; setting it aside leaves 7 of 9 on the ResNet-18 features. Against the whole set of frozen encoders it lands just below: the strongest encoder for each cohort averages 0.409, and TRIPLEX is above that cohort’s best encoder in one of the ten (CCRCC). A method designed for multi-resolution reading therefore improves on a linear head, and still scores at the level the scalar rewards.

**Table 9:** TRIPLEX against ridge on the same features, per cohort. Folds are the benchmark’s own. TRIPLEX entries are the mean over folds of its best checkpoint on that fold, its authors’ rule; ridge uses a penalty chosen on the other nine cohorts. The last column is the best of the fifty-seven frozen encoders for that cohort. On IDC the ResNet-18 run completed 1 of the 4 folds, and its entry († ) is that fold’s value.

| Cohort | Folds | CIGA (ResNet-18) features |  | UNI features |  | Best of 57 |
| --- | --- | --- | --- | --- | --- | --- |
|  |  | TRIPLEX | ridge | TRIPLEX | ridge | ridge |
| SKCM | 2 | 0.601 | 0.430 | 0.642 | 0.614 | 0.685 |
| HCC | 2 | 0.085 | 0.057 | 0.106 | 0.085 | 0.110 |
| LUNG | 2 | 0.582 | 0.534 | 0.587 | 0.587 | 0.622 |
| PAAD | 3 | 0.449 | 0.400 | 0.459 | 0.456 | 0.500 |
| COAD | 2 | 0.309 | 0.329 | 0.312 | 0.344 | 0.403 |
| READ | 2 | 0.121 | 0.098 | 0.249 | 0.244 | 0.282 |
| IDC | 4 | 0.462† | 0.429 | 0.577 | 0.574 | 0.597 |
| LYMPH.IDC | 4 | 0.260 | 0.269 | 0.272 | 0.294 | 0.306 |
| PRAD | 2 | 0.295 | 0.172 | 0.317 | 0.318 | 0.366 |
| CCRCC | 6 | 0.220 | 0.156 | 0.254 | 0.199 | 0.222 |
| Mean | | 0.339† | 0.288 | 0.377 | 0.372 | 0.409 |

Two of the 72 slides needed a departure from the authors’ pipeline, both in building the neighbourhood view. For NCBI684 the tissue mask left only a quarter of the 1120-pixel windows, so we extracted that slide’s neighbourhoods without the mask; the other seventy keep every window under the mask. For TENX111 the coordinate frame in the raw release does not match the benchmark’s own patches, and every window fell outside the image, so we cut its neighbourhoods from the slide at the benchmark’s coordinates, which we checked against the stored patches. Both slides sit in the two cohorts where TRIPLEX trails ridge on the ResNet-18 features.

HECLIP has no projection head on the expression side. Its reference embedding is the expression vector itself, so the projection dimension must equal the number of genes. The authors’ released configurations use panels of roughly 3400 genes. This benchmark scores 50 genes. HECLIP is therefore run far outside the regime it was designed for.

HGGEP builds a hypergraph by looping over spots in Python, once per forward pass. At the 4000 spots this protocol allows, that loop issues about twenty thousand kernel calls and dominates the runtime. We replaced it with a vectorised form. We verified that form against the authors’ original at three sizes. The edge sets are bit-identical and the weights agree to 9.3 × 10^*−*10^. The model also needs a large accelerator at this spot count, and we ran it on 180 GB GPUs.

THItoGene (Jia et al., 2024) shares its data pipeline with Hist2ST, and we ran it through the same adaptor. One line of the authors’ capsule layer divides a GPU tensor by a constant built on the CPU. Current PyTorch rejects that. We replaced that constant with a scalar of the same value. The method is also the least stable of the six from one run to the next. On the first SKCM fold, two runs differed only in their random initialisation. They scored 0.27 and 0.10. Table 8 reports the first full run, as for the other five. Two further full runs with fixed seeds score 0.0747 and 0.0715 against the first run’s 0.0519 (mean 0.066 ± 0.012). Each of the three sits below every one of the 57 encoders.

## I The full benchmark table

Table 10 summarises Table 1 by encoder size, and Table 11 lists every encoder with both scores and the paired differences, ordered by the trained model’s own score. The size correlations of Section 3.2 rank each encoder’s oracle-to-model ratio (Table 1) against its parameter count, with two-sided permutation *P* values from 20,000 permutations. A family merges encoders that share a training recipe or a teacher and takes its members’ medians, which gives 32 families, 29 without the three made up only of distilled students. Spearman *r*_*s*_ is −0.30 over encoders (*P* = 0.024) and −0.37 over families (*P* = 0.035); without the distilled encoders it is −0.60 over 49 encoders (*P <* 10^*−*4^) and −0.58 over 29 families (*P* = 0.0013).

**Table 10:**
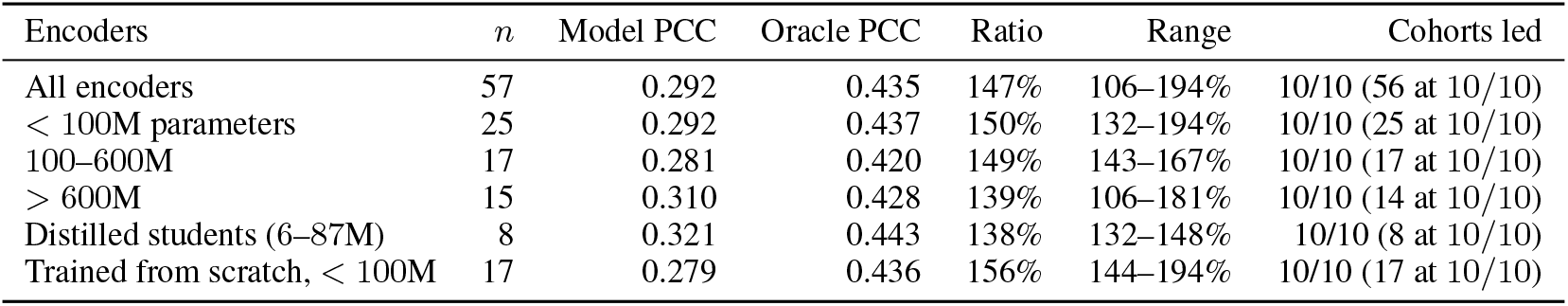
The domain oracle against a trained model on the public benchmark (72 samples, 10 cohorts, 57 frozen encoders, *K* = 20), summarised by encoder size. Model and Oracle are medians over encoders of the mean PCC over samples; Ratio is the median of Oracle over Model and Range its extremes; Cohorts led is the median number of cohorts in which the oracle leads, with the number of encoders whose oracle leads in all ten. Table 11 lists all 57 encoders

| Encoders | $n$ | Model PCC | Oracle PCC | Ratio | Range | Cohorts led |
| --- | --- | --- | --- | --- | --- | --- |
| All encoders | 57 | 0.292 | 0.435 | 147% | 106–194% | 10/10 (56 at 10/10) |
| < 100M parameters | 25 | 0.292 | 0.437 | 150% | 132–194% | 10/10 (25 at 10/10) |
| 100–600M | 17 | 0.281 | 0.420 | 149% | 143–167% | 10/10 (17 at 10/10) |
| > 600M | 15 | 0.310 | 0.428 | 139% | 106–181% | 10/10 (14 at 10/10) |
| Distilled students (6–87M) | 8 | 0.321 | 0.443 | 138% | 132–148% | 10/10 (8 at 10/10) |
| Trained from scratch, < 100M | 17 | 0.279 | 0.436 | 156% | 144–194% | 10/10 (17 at 10/10) |

**Table 11:**
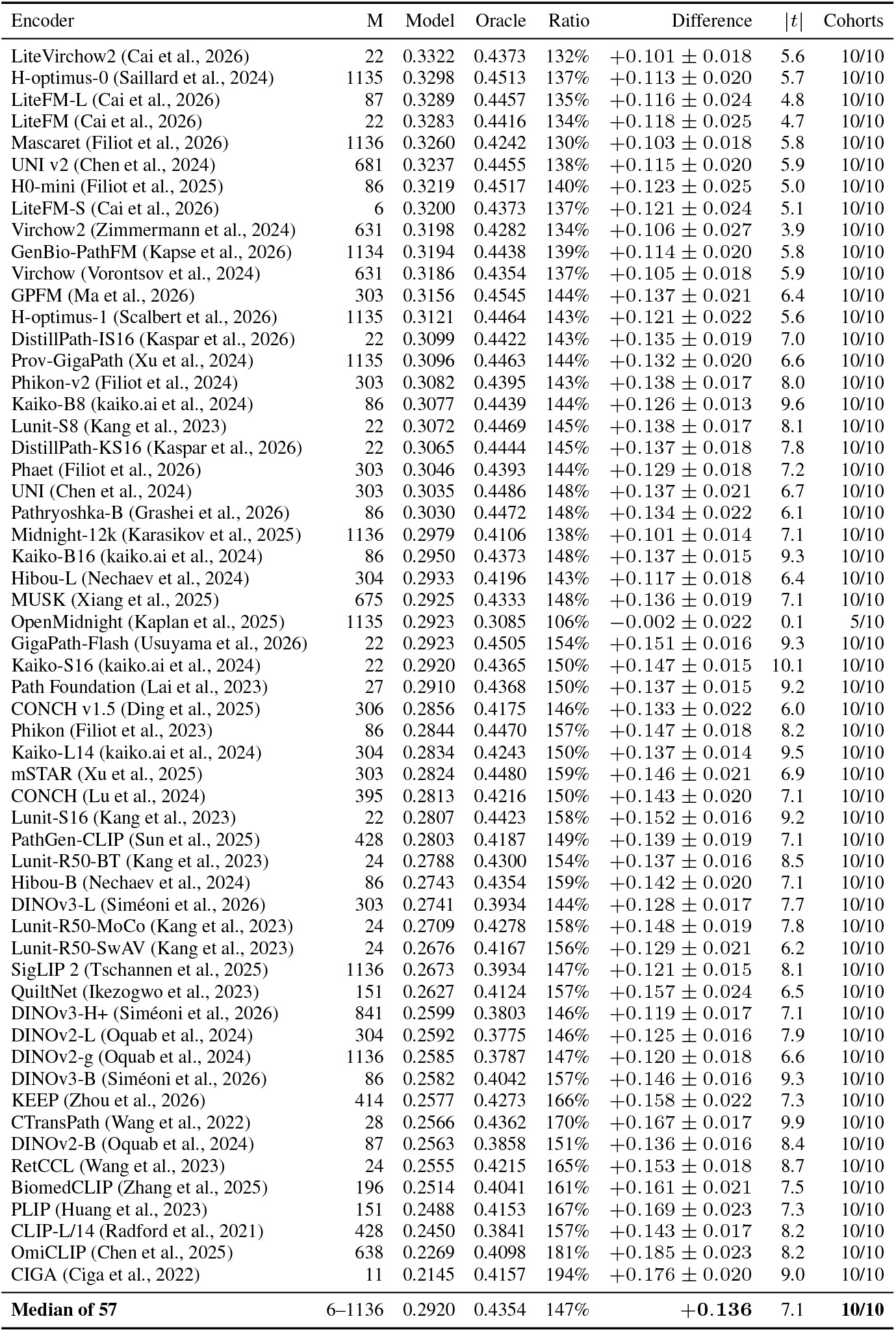
The domain oracle of Proposition 1 against a trained model, on 72 samples and 10 cohorts, for each of 57 frozen encoders spanning 6M to 1.1B parameters, ordered by the model’s own score. Oracle and Model are mean PCC over samples; Ratio is Oracle over Model; M is the parameter count in millions, measured on the loaded encoder; Difference is Oracle minus Model in PCC units, as the mean and its standard error over the ten per-cohort median paired differences; |*t*| is that mean over its standard error; Cohorts counts those in which the oracle leads. *K* = 20. Table 10 summarises these rows; the one encoder whose oracle does not lead in all ten cohorts, OpenMidnight, is explained in Appendix J

## J Feature standardisation and the OpenMidnight exception

OpenMidnight is the only encoder in Table 11 whose domain oracle does not beat the trained model in the cohort-level paired comparison at *K* = 20: it leads in 5 of 10 cohorts with a mean paired difference of −0.002, although its mean over samples is 6% higher. Its model score is unremarkable, 0.29 against a median of 0.29. Its oracle is not. It scores 0.31 against a median of 0.44. Midnight-12k is the same size, and OpenMidnight reproduces its recipe. Yet OpenMidnight’s oracle is below Midnight-12k’s on all 72 samples. At *K* = 50 its oracle leads the trained model in 9 of 10 cohorts and at *K* = 200 in all 10, so the anomaly lives in the coarse partition and not in what the features carry.

The cause is a single coordinate. We looked at six samples spanning five cohorts. One of OpenMidnight’s 1536 feature dimensions carries 0.88 to 0.94 of the total variance, and the first principal component 92% on average. We compared four other encoders. There the largest single dimension carries at most 0.06 and the first component 20% to 33%. Our partition is *K*-means on the first 50 principal components of the raw features, so at *K* = 20 it splits mostly along that one coordinate. The resulting clusters are less spatially coherent than the others’: 0.19 of a spot’s six nearest neighbours share its cluster, against 0.33 to 0.45. The coordinate is not brightness. Its correlation with the patch’s mean intensity is 0.34. That is within the range of the same correlation for the other encoders. Outlier tokens with anomalous norms are a known artefact of large vision transformers (Darcet et al., 2024); what we see here is an outlier coordinate of the pooled embedding.

Standardising each feature dimension to unit variance before the projection removes the effect. Table 12 repeats the *K* = 20 oracle for all 57 encoders under that one change. OpenMidnight’s oracle rises from 0.31 to 0.42, its ratio from 106% to 145%, and it wins 10 of 10 cohorts. No other encoder’s oracle moves by more than 0.009, no other ratio by more than three points, and no other cohort count changes. We keep the unstandardised recipe in the main text because it is the one we fixed before seeing the results. Under the unstandardised recipe the bound of Proposition 1 sits above the trained model in the mean over samples for all 57 and cohort by cohort for 56; under the standardised recipe it does so for all 57 on both readings.

**Table 12:** The domain oracle at *K* = 20 with the features standardised before the projection (std.), against the recipe of Table 11, for all 57 encoders, ordered by the change. Oracle is mean PCC over samples; Difference is Oracle, std. minus Oracle; Ratio is Oracle over the same trained model; and Cohorts counts those the oracle wins under the two recipes; the last row is the median of each column over the 57

| Encoder | Oracle | Oracle, std. | Difference | Ratio | Ratio, std. | Cohorts, raw → std. |
| --- | --- | --- | --- | --- | --- | --- |
| H0-mini (Filiot et al., 2025) | 0.452 | 0.450 | -0.002 | 140% | 140% | 10→10 |
| DINOv3-B (Siméoni et al., 2026) | 0.404 | 0.403 | -0.002 | 157% | 156% | 10→10 |
| GigaPath-Flash (Usuyama et al., 2026) | 0.451 | 0.449 | -0.001 | 154% | 154% | 10→10 |
| KEEP (Zhou et al., 2026) | 0.427 | 0.426 | -0.001 | 166% | 165% | 10→10 |
| Lunit-R50-MoCo (Kang et al., 2023) | 0.428 | 0.426 | -0.001 | 158% | 157% | 10→10 |
| Phaet (Filiot et al., 2026) | 0.439 | 0.438 | -0.001 | 144% | 144% | 10→10 |
| Lunit-S8 (Kang et al., 2023) | 0.447 | 0.446 | -0.001 | 145% | 145% | 10→10 |
| DistillPath-IS16 (Kaspar et al., 2026) | 0.442 | 0.441 | -0.001 | 143% | 142% | 10→10 |
| Kaiko-S16 (kaiko.ai et al., 2024) | 0.437 | 0.436 | -0.001 | 150% | 149% | 10→10 |
| DistillPath-KS16 (Kaspar et al., 2026) | 0.444 | 0.444 | -0.001 | 145% | 145% | 10→10 |
| UNI (Chen et al., 2024) | 0.449 | 0.448 | -0.001 | 148% | 148% | 10→10 |
| CONCH (Lu et al., 2024) | 0.422 | 0.421 | -0.000 | 150% | 150% | 10→10 |
| Kaiko-B16 (kaiko.ai et al., 2024) | 0.437 | 0.437 | -0.000 | 148% | 148% | 10→10 |
| Lunit-S16 (Kang et al., 2023) | 0.442 | 0.442 | -0.000 | 158% | 157% | 10→10 |
| OmiCLIP (Chen et al., 2025) | 0.410 | 0.410 | -0.000 | 181% | 181% | 10→10 |
| CONCH v1.5 (Ding et al., 2025) | 0.418 | 0.418 | -0.000 | 146% | 146% | 10→10 |
| Phikon (Filiot et al., 2023) | 0.447 | 0.447 | +0.000 | 157% | 157% | 10→10 |
| BiomedCLIP (Zhang et al., 2025) | 0.404 | 0.404 | +0.000 | 161% | 161% | 10→10 |
| Kaiko-B8 (kaiko.ai et al., 2024) | 0.444 | 0.444 | +0.000 | 144% | 144% | 10→10 |
| Hibou-B (Nechaev et al., 2024) | 0.435 | 0.436 | +0.000 | 159% | 159% | 10→10 |
| Lunit-R50-BT (Kang et al., 2023) | 0.430 | 0.430 | +0.001 | 154% | 154% | 10→10 |
| DINOv2-B (Oquab et al., 2024) | 0.386 | 0.386 | +0.001 | 151% | 151% | 10→10 |
| CTransPath (Wang et al., 2022) | 0.436 | 0.437 | +0.001 | 170% | 170% | 10→10 |
| GPFM (Ma et al., 2026) | 0.454 | 0.455 | +0.001 | 144% | 144% | 10→10 |
| Pathryoshka-B (Grashei et al., 2026) | 0.447 | 0.448 | +0.001 | 148% | 148% | 10→10 |
| DINOv3-L (Siméoni et al., 2026) | 0.393 | 0.395 | +0.001 | 144% | 144% | 10→10 |
| PathGen-CLIP (Sun et al., 2025) | 0.419 | 0.420 | +0.001 | 149% | 150% | 10→10 |
| Mascaret (Filiot et al., 2026) | 0.424 | 0.426 | +0.001 | 130% | 131% | 10→10 |
| GenBio-PathFM (Kapse et al., 2026) | 0.444 | 0.445 | +0.002 | 139% | 139% | 10→10 |
| PLIP (Huang et al., 2023) | 0.415 | 0.417 | +0.002 | 167% | 168% | 10→10 |
| Phikon-v2 (Filiot et al., 2024) | 0.440 | 0.441 | +0.002 | 143% | 143% | 10→10 |
| Prov-GigaPath (Xu et al., 2024) | 0.446 | 0.448 | +0.002 | 144% | 145% | 10→10 |
| DINOv3-H+ (Siméoni et al., 2026) | 0.380 | 0.382 | +0.002 | 146% | 147% | 10→10 |
| DINOv2-g (Oquab et al., 2024) | 0.379 | 0.381 | +0.002 | 147% | 147% | 10→10 |
| mSTAR (Xu et al., 2025) | 0.448 | 0.450 | +0.002 | 159% | 159% | 10→10 |
| SigLIP 2 (Tschannen et al., 2025) | 0.393 | 0.395 | +0.002 | 147% | 148% | 10→10 |
| UNI v2 (Chen et al., 2024) | 0.445 | 0.448 | +0.002 | 138% | 138% | 10→10 |
| Path Foundation (Lai et al., 2023) | 0.437 | 0.439 | +0.002 | 150% | 151% | 10→10 |
| H-optimus-1 (Scalbert et al., 2026) | 0.446 | 0.449 | +0.002 | 143% | 144% | 10→10 |
| DINOv2-L (Oquab et al., 2024) | 0.378 | 0.380 | +0.002 | 146% | 147% | 10→10 |
| H-optimus-0 (Saillard et al., 2024) | 0.451 | 0.454 | +0.003 | 137% | 138% | 10→10 |
| Kaiko-L14 (kaiko.ai et al., 2024) | 0.424 | 0.427 | +0.003 | 150% | 151% | 10→10 |
| RetCCL (Wang et al., 2023) | 0.421 | 0.425 | +0.004 | 165% | 167% | 10→10 |
| Midnight-12k (Karasikov et al., 2025) | 0.411 | 0.415 | +0.004 | 138% | 139% | 10→10 |
| QuiltNet (Ikezogwo et al., 2023) | 0.412 | 0.416 | +0.004 | 157% | 159% | 10→10 |
| CIGA (Ciga et al., 2022) | 0.416 | 0.420 | +0.004 | 194% | 196% | 10→10 |
| CLIP-L/14 (Radford et al., 2021) | 0.384 | 0.389 | +0.005 | 157% | 159% | 10→10 |
| MUSK (Xiang et al., 2025) | 0.433 | 0.438 | +0.005 | 148% | 150% | 10→10 |
| LiteVirchow2 (Cai et al., 2026) | 0.437 | 0.443 | +0.005 | 132% | 133% | 10→10 |
| LiteFM (Cai et al., 2026) | 0.442 | 0.447 | +0.005 | 134% | 136% | 10→10 |
| LiteFM-S (Cai et al., 2026) | 0.437 | 0.443 | +0.006 | 137% | 138% | 10→10 |
| LiteFM-L (Cai et al., 2026) | 0.446 | 0.452 | +0.006 | 135% | 137% | 10→10 |
| Lunit-R50-SwAV (Kang et al., 2023) | 0.417 | 0.424 | +0.007 | 156% | 158% | 10→10 |
| Virchow (Vorontsov et al., 2024) | 0.435 | 0.442 | +0.007 | 137% | 139% | 10→10 |
| Hibou-L (Nechaev et al., 2024) | 0.420 | 0.427 | +0.008 | 143% | 146% | 10→10 |
| Virchow2 (Zimmermann et al., 2024) | 0.428 | 0.437 | +0.008 | 134% | 137% | 10→10 |
| OpenMidnight (Kaplan et al., 2025) | 0.309 | 0.423 | +0.114 | 106% | 145% | 5→10 |
| **Median of 57** | 0.435 | 0.437 |  | 147% | 148% |  |

## K Additional results

### K.1 The scalar score tracks tissue autocorrelation

If a scalar correlation mostly rewards coarse spatial structure, it should track how strongly structured the tissue is rather than how good the model is. It does.

Predicted fields are systematically smoother than measured ones. On the sixteen regions of the first release, Moran’s *I* (Moran, 1950) *rises from measurement to prediction in 16/*16 regions and 8*/*8 specimens, with median gap +0.32 and specimen range 0.18 to 0.51 (exact *P* = 0.0078). The inflation carries information beyond the score. It moves with the effective resolution at *r* = +0.912 across specimen medians, and the partial correlation is +0.826 given PCC and +0.513 given the noise ceiling.

Across regions the reported score tracks the measured tissue’s own spatial autocorrelation, with *r* = 0.828 and *R*^2^ = 0.686 (specimen-level permutation *P* = 0.0076). The same relation holds within regions, gene by gene, in the fifteen regions (from seven specimens) that have per-gene calibration curves. Genes whose measured field is more autocorrelated are predicted better, and that confound falls on the scalar much harder than on the effective resolution *σ*_A_. The across-gene correlation with Moran’s *I* has specimen median +0.85 for PCC and −0.40 for log *σ*. Here *σ* is read off a per-gene calibration curve, that is, (4) computed on a single gene. The difference between the two correlations holds in 7*/*7 specimens (*P* = 0.0156).

Measurement noise does not explain the pattern. A split-half noise ceiling has median 0.93 and exceeds the achieved score in 16*/*16 regions, so a real gap remains after noise is accounted for. Dividing the score by that ceiling barely reorders the regions, with Spearman *r*_*s*_ = 0.926 and a maximum shift of four places. Figure 4 collects these results in six panels.

**Figure 4.**
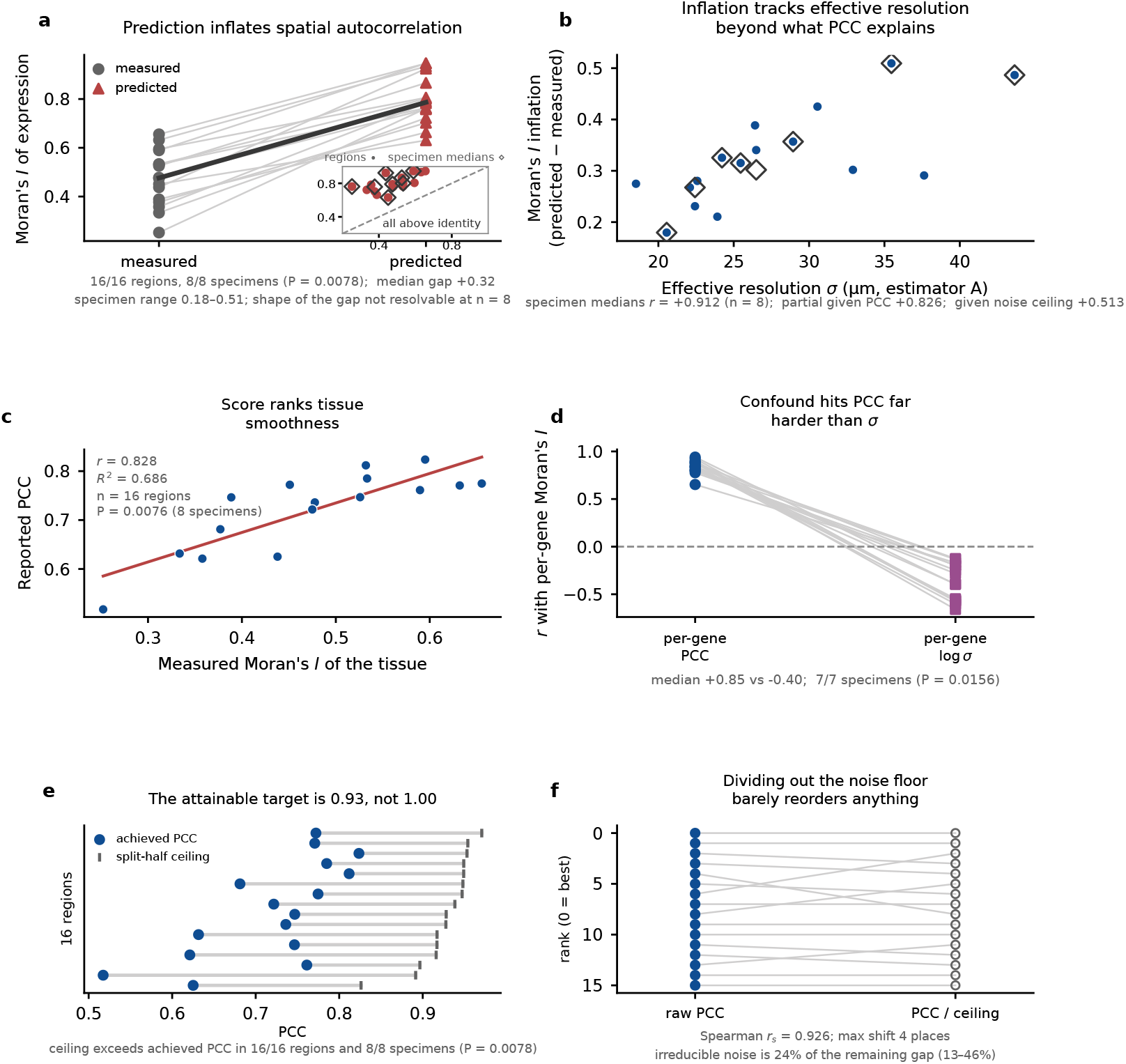
The quantities the reported score tracks. **(a)** Moran’s *I*, measured against predicted. **(b)** Its inflation against effective resolution. **(c)** Reported score against the tissue’s own autocorrelation. **(d)** The same relation gene by gene, for the score and for the effective resolution *σ*_A_. **(e–f)** A split-half noise ceiling, and ranks before and after dividing by it. Inference is at the specimen level throughout. This analysis uses the sixteen regions of the first release, and panel d the fifteen of them with per-gene calibration curves

### K.2 Unreported protocol choices

The gene-panel and patch-size sweeps below run on the two Visium HD colon sections with one frozen encoder; the encoder span they are compared against is taken across twenty-five encoders on the same two sections. The normalisation result is on the benchmark, over all 57.

One choice that papers rarely state moves the score by more than the entire spread across encoders. The genes scored are the ones with highest variance. Raising their number from 20 to 200 moves the score by 0.197, where the span of all twenty-five encoders under the same protocol is only 0.116. The gene panel does this by changing the target itself. A second choice moves the score by two thirds of the encoder span on the benchmark: changing the expression normalisation from log(1 + *x*) to counts-per-10^4^ followed by log(1 + *x*) lowers the reported score by 31.0%, a move of 0.079 against the encoders’ span of 0.121 on the benchmark, in 57*/*57 encoders and 10*/*10 cohorts. That change shifts the magnitude and largely keeps the ranking: Spearman *r*_*s*_ = 0.975, with 86 discordant pairs out of 1596. The median encoder moves two places, nine move five or more, and one moves fifteen.

The two choices of Figure 5d, the gene panel and the encoder patch size, do not act on the same quantity. The gene panel moves the score. The encoder patch size, from 224 to 1344 pixels, moves the score by less but the effective resolution *σ*_A_ by more. It leaves the target and its calibration curve numerically unchanged. So under the two choices the same change in score corresponds to very different changes in blur.

**Figure 5.**
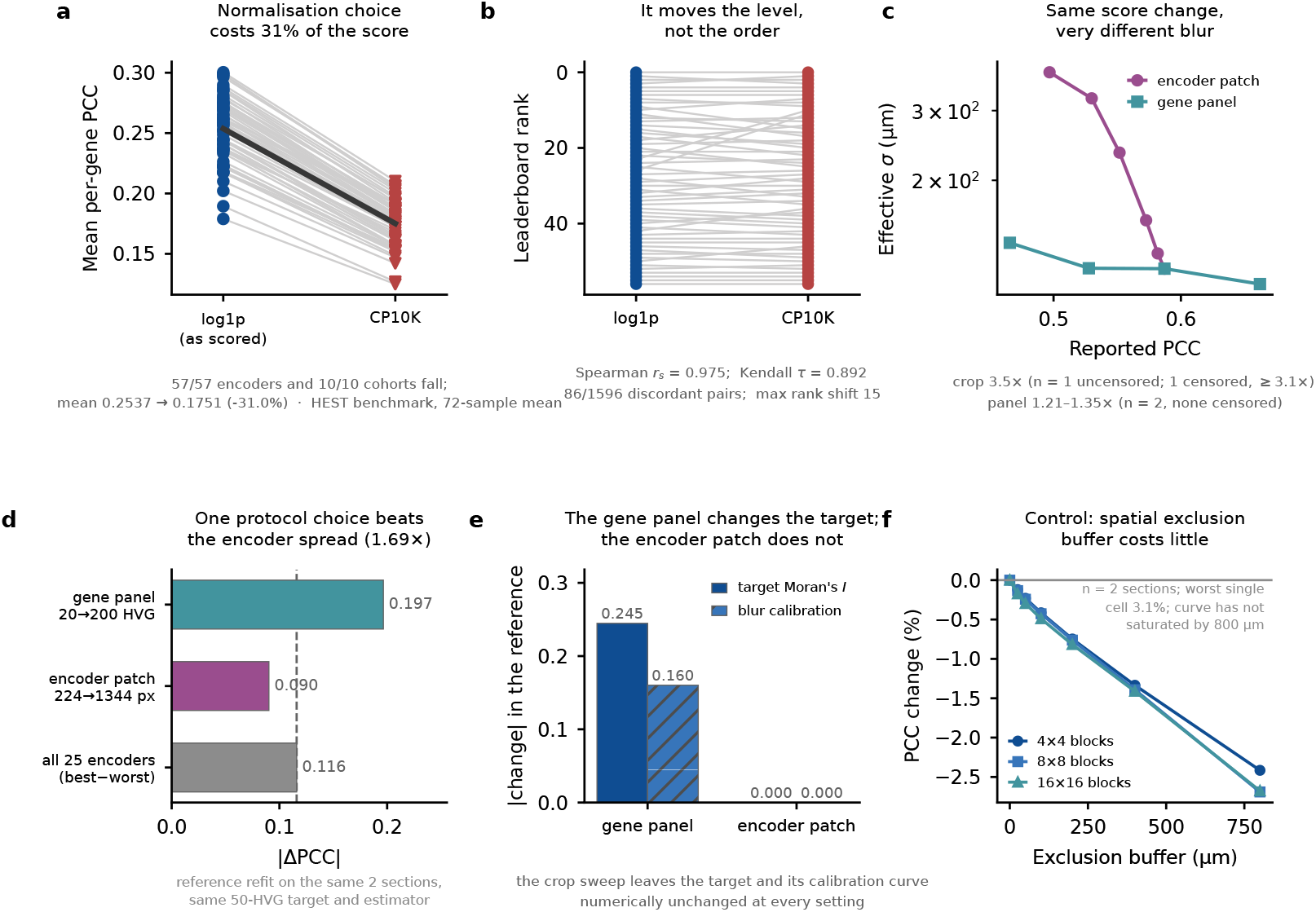
Unreported choices in the evaluation protocol. **(a–b)** The expression normalisation, on the score and on the ranking. **(c)** What a given change in score corresponds to in blur. **(d)** Each protocol choice against the spread across 25 encoders on the two Visium HD sections. **(e)** The gene panel changes the target and its calibration curve; the encoder patch leaves both unchanged. **(f)** A control on the spatial exclusion buffer

The large movements above are specific to these protocol choices. Widening the spatial exclusion buffer to 800 *µ*m changes the score by at most 3.1% in any single section and block grid, though the score-versus-buffer curve has not flattened by 800 *µ*m. Figure 5 collects the results of this subsection in six panels.

### K.3 Band-wise reliability

Counting noise enters the finest bands first, so a ceiling on the scalar says little about *β*_1_. You et al. (2024) document across platforms how spatial data quality depends on sequencing depth and capture sensitivity. For each region we split every bin’s counts at random into two halves by binomial thinning, apply the same normalisation and the same band decomposition to both halves, and correlate the two halves band by band and gene by gene. Thinning models counting noise only; optical crowding, off-target probes and segmentation errors are shared by the two halves, so the ceiling does not account for them. The mean over genes is the split-half reliability *c*_1*/*2_. Its Spearman–Brown extrapolation to the full depth is *c* = 2*c*_1*/*2_*/*(1 + *c*_1*/*2_). Then 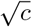 bounds the correlation of a noiseless predictor with the measured band. Table 13 gives medians and ranges over the 22 regions, three random splits each. The finest band is the least reliable, *c* = 0.59 with a range of 0.30 to 0.71 and the four lowest values all on Xenium Prime regions; and reliability rises with width to 0.98 at the coarsest scale.

**Table 13:**
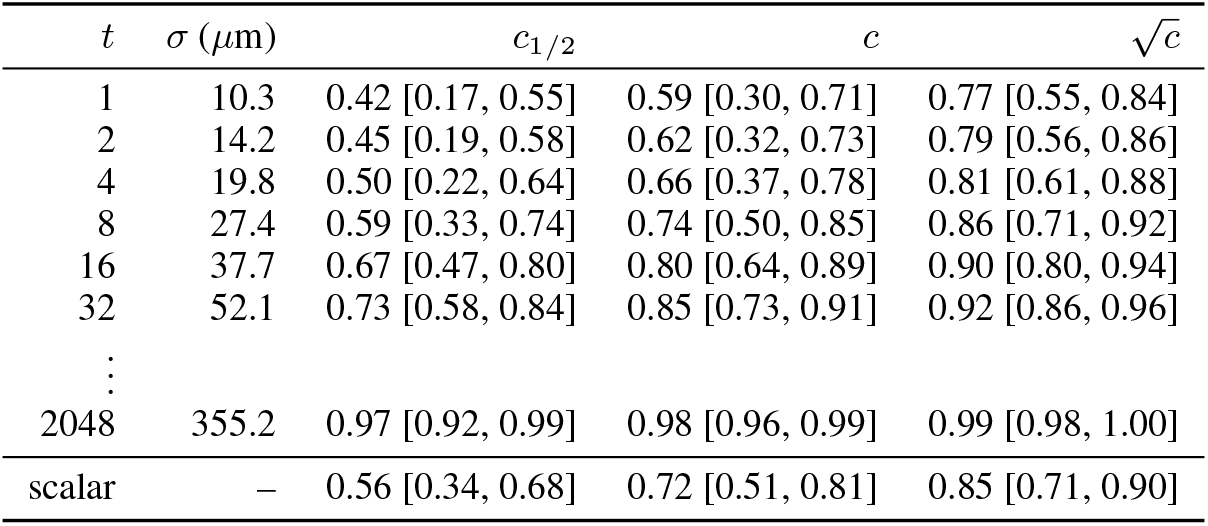
Split-half reliability by band on the 22 Xenium regions, median [min, max]. *c*_1*/*2_ is the correlation between two binomial halves of the counts after band filtering, *c* its Spearman–Brown extrapolation to the full depth, and 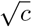 the ceiling on any predictor’s correlation with the measured band. The last row is the unfiltered field.

Δ_fine_ is insensitive to counting noise. Take a predictor independent of the measurement noise. Attenuation multiplies its *β*_1_ by 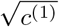, where *c*^(1)^ is the finest band’s reliability from Table 13, and a normalised difference of two such values cancels it. The oracle is built from the measured field, so it is not independent. Appendix K.4 therefore checks the claim directly with a cross-fitted oracle. It finds the ratio Δ_fine_*/*Δ_scalar_ unchanged to one decimal place. The attenuation-corrected levels, formed per region before the median, are 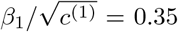 for the model and 0.06 for the oracle.

The finest-band variance share of the measured field in Section 3.3 contains noise. We keep only the reproducible part. The band’s share of reproducible variance is then *c*^(1)^*/c* times the raw share, where *c* is the reliability of the unfiltered field. The median falls from 19% to 15% (range 9–20%).

Against that share both predictors are still too smooth by a factor of 2.3 to 7.2 (median 4.3), rather than the 2.6 to 12.3 of the raw share. Each factor is formed per region before the medians, so the median factor is not the quotient of the median shares. On the 200-gene panel of that section the scalar ceiling is 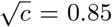. The 0.93 of Appendix K.1 is the same estimator on a 50-gene panel whose genes are more highly expressed.

### K.4 Partition controls and known degradations

Tables 14 and 15 test, on the benchmark and on Xenium, whether the partition needs the image and whether a predictor allowed nothing but domain identity can be learned rather than handed the answer. Each control keeps the oracle’s recipe and changes one ingredient. A *coordinate partition* replaces the image features by the spot coordinates in the *K*-means step, so its clusters are spatial blocks that know nothing about the tissue. A *random matched-size partition* permutes the image partition’s labels across spots. This keeps the cluster sizes and destroys everything else. Its score is the finite-sample optimism of filling *K* clusters with measured means. The *train-only* variants keep a partition and estimate each cluster’s mean from training data alone. On the benchmark that data is the other samples of the same cohort, assigned to the test sample’s clusters through that sample’s own principal components and centroids. On Xenium it is the training blocks of the same spatial fold. These are predictors, not bounds. The image oracle in both tables is the value reported in the main text; recomputing it alongside the controls moves the per-encoder median by 0.001 at the median over encoders and by at most 0.008, while single samples move by up to 0.054. That per-sample spread is larger than the third-decimal restart variation that Appendix E measures on Xenium; it comes from the partition landing in a different local optimum on individual samples. The *inductive* variant goes one step further and learns the partition as well: the PCA and the *K*-means are fitted on training data only and each test spot is assigned to its nearest centre, so nothing about the test sample enters the predictor but its own image features. It scores 0.204 on the benchmark, 79% of the ridge model (range 41–92% over encoders, against 82% with the partition fitted on the test sample), and 0.423 on Xenium, 78% against 79%. Where the partition is fitted does not matter; what it is fitted to does.

**Table 14:** Partition controls on the benchmark, 57 encoders. Entries are the median over encoders of each encoder’s median PCC over the 72 samples (Table 10 uses each encoder’s mean over samples instead, hence its higher levels); the coordinate partitions do not depend on the encoder, and the random partition depends on it only through the cluster sizes it inherits. The last column is the median over encoders of the number of cohorts in which the control’s median exceeds the ridge model’s.

| Predictor on the benchmark | PCC (median over encoders) | Range over encoders | Cohorts above ridge |
| --- | --- | --- | --- |
| Ridge on frozen features (cohort-selected $\lambda$) | 0.268 | 0.181–0.329 | – |
| Image partition, measured means (domain oracle) | 0.414 | 0.285–0.442 | – |
| Coordinate partition, measured means | 0.361 | 0.361–0.361 | 6/10 |
| Random matched-size partition, measured means | 0.080 | 0.076–0.084 | – |
| Image partition, means from training samples | 0.246 | 0.150–0.274 | 2/10 |
| Coordinate partition, means from training samples | 0.058 | 0.058–0.058 | 0/10 |
| Image partition learned on training samples, means from training samples | 0.204 | 0.079–0.267 | 2/10 |

**Table 15:** Partition controls on Xenium. PCC is the median over specimens of the per-specimen median over regions; the last column is the same aggregation of the per-region ratio to the ridge model, so the image oracle’s 0.79 is the 79% of Table 2.

| Predictor on Xenium (22 regions, 13 specimens) | PCC | Range over specimens | Fraction of ridge |
| --- | --- | --- | --- |
| Ridge on frozen features | 0.559 | 0.423–0.672 | 1.00 |
| Image partition, measured means ($K = 20$; domain oracle) | 0.428 | 0.292–0.600 | 0.79 |
| Coordinate partition, measured means ($K = 20$) | 0.186 | 0.151–0.459 | 0.33 |
| Random matched-size partition, measured means ($K = 20$) | 0.011 | 0.007–0.021 | 0.02 |
| Image partition, means from training blocks ($K = 20$) | 0.427 | 0.289–0.599 | 0.79 |
| Image partition learned on training blocks, means from training blocks ($K = 20$) | 0.423 | 0.287–0.589 | 0.78 |
| Coordinate partition, means from training blocks ($K = 20$) | 0.159 | 0.117–0.441 | 0.29 |
| Image partition, measured means ($K = 200$) | 0.471 | 0.332–0.629 | 0.88 |
| Coordinate partition, measured means ($K = 200$) | 0.293 | 0.234–0.536 | 0.57 |
| Image partition, means from training blocks ($K = 200$) | 0.467 | 0.326–0.626 | 0.86 |

**Table 16:** Known degradations of the measured field and their finest-band correlation. The first three are tuned to the trained model’s scalar score; hotspot removal cannot reach it even at its strongest setting. Medians over specimens of per-specimen medians over regions.

| Field compared with the measurement | Scalar PCC | $\beta_1$ | $\beta_1$ range over specimens |
| --- | --- | --- | --- |
| Trained model (ridge) | 0.559 | 0.238 | 0.124–0.354 |
| Measurement blurred (diffusion, matched) | 0.563 | 0.088 | 0.042–0.205 |
| Measurement shifted (boundary displacement, matched) | 0.563 | 0.179 | 0.103–0.234 |
| Measurement plus white noise (matched) | 0.559 | 0.474 | 0.381–0.548 |
| Measurement with hotspots removed (strongest setting, not matchable) | 0.896 | 0.855 | 0.825–0.883 |
| Domain oracle ($K = 20$) | 0.428 | 0.039 | 0.010–0.170 |

On the benchmark the levels below are medians over samples, then the median over the 57 encoders, while every benchmark percentage in this paragraph is a median over encoders of the per-cohort ratio, and every Xenium percentage a median over specimens of the per-region ratio, so none of them is the quotient of the levels we print. The trained model scores 0.268, the domain oracle 0.414, the coordinate partition 0.361, the random partition 0.080 and the train-only image partition 0.246. A coordinate partition with means from training samples scores 0.058, above the model in 0/10 cohorts: a position on the slide carries nothing from one sample to the next. A predictor allowed only the identity of 20 image-derived domains, with means learned from other samples, still reaches a median of 82% of the trained model’s score (range 61–92% over encoders), down from 140% with measured means, so the mechanism the oracle isolates is learnable. At the benchmark’s 100 *µ*m pitch the partition does not need the image: 20 spatial blocks with measured means reach 80% of the image oracle and 123% of the model, above the model in a median 6 of 10 cohorts. The random partition puts the optimism of measured means at 0.080, 17% of the image oracle. On Xenium at 16 *µ*m the image does matter. Coordinate blocks reach 33% of the model against the image oracle’s 79%. The random partition reaches 2% and the train-only image partition 79%. That last value is within 1% of the oracle in all 13 specimens, because the training blocks of a section carry the same domains as its test blocks. At *K* = 200 the same ordering holds at higher levels. On the two Visium HD colon sections the same oracle at *K* = 20 reaches 88% to 94% of the ridge model over the four within-slide folds. The image partition is thus one coarse partition among others on the benchmark and the informative one at 16 *µ*m; in both settings what the score rewards is the identity of a coarse partition, not the structure inside its clusters.

Table 16 asks the converse question: does the finest band separate degradations that the scalar cannot? We degrade the measured field itself in four known ways and tune the first three until their correlation with the measurement equals the trained model’s score in each region, to within 0.004, then read *β*_1_. Diffusion blur, boundary displacement (the field shifted by a fixed distance along one axis) and additive white noise all reach the model’s scalar score. At that point their finest-band correlations are 0.09, 0.18 and 0.47. The model itself sits at 0.24. It is above the blurred and the shifted field and below the noisy one. Removing hotspots replaces each gene’s top values by a locally smoothed value. Even at its strongest setting (top 60% of values replaced, PCC 0.90) it cannot be pushed down to the model’s score. Its finest band stays at 0.85. One scalar value therefore covers finest-band fidelities from 0.09 to 0.47.

### The model’s score sits in its between-cluster part, and its finest-band fidelity in the within-cluster part

The oracle bounds what a partition can achieve. To see what the trained model uses, Table 17 splits the model’s own prediction into its between-cluster part **Π**_*P*_ **Ŷ** and its within-cluster part (**I™ Π**_*P*_ ) **Ŷ** on the same image partition. The split uses no test label. The between-cluster part alone scores 0.428 [0.291, 0.600] against the measurement. That is a median 79% of the model’s own score, the ratio formed per region (range 69–89%). The between-cluster part also equals the oracle’s score to three decimals. The model’s cluster means track the measured ones almost perfectly. The within-cluster part correlates 0.377 [0.297, 0.460] with the within-cluster measurement, so the model does recover structure inside clusters, and that part holds 38% of the model’s variance against 80% of the measurement’s. In the finest band the separation is clean: the between-cluster part has *β*_1_ = 0.04, the oracle’s value, while the within-cluster part has 0.29 and the full model 0.24. Everything the finest band credits to the model is within-cluster structure, and everything the scalar credits to the oracle is between-cluster structure. The 79% ratio is not an additive share, since the two correlations carry different denominators. With a common denominator the per-gene correlation splits exactly into a between-cluster and a within-cluster term, *ρ*(*ŷ, y*) = ⟨**Π***ŷ*, **Π***y*⟩ */*( ∥*ŷ*∥ ∥*y*∥ ) + ⟨ (**I** − **Π**)*ŷ*, (**I** − **Π**)*y*⟩ */*( ∥*ŷ*∥ ∥*y*∥ ) for centred vectors, and on the 22 regions the between-cluster term accounts for a median 64% of the model’s score (range 49–80% over specimens) and the within-cluster term for 36%.

**Table 17:**
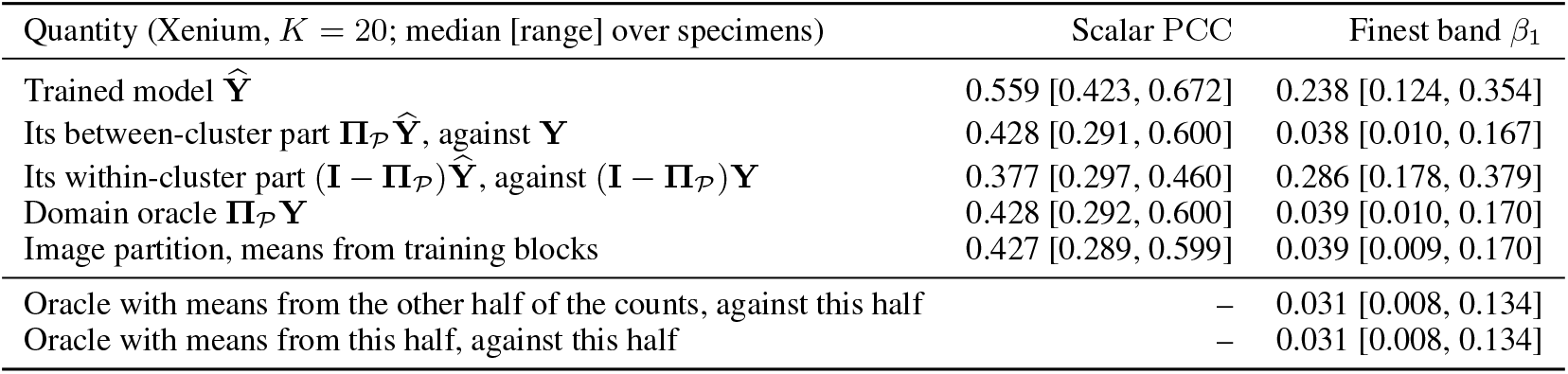
The trained model split on the image partition, the oracle, a learnable partition predictor, and the cross-fitted oracle, on the Xenium regions. Medians over specimens of per-specimen medians over regions; the cross-fitted rows use two binomial halves of the counts.

#### Cross-fitting the oracle leaves its finest band unchanged, and a learned domain predictor gives the same result

To test the attenuation argument of Appendix K.3 on the oracle, we split each bin’s counts into two binomial halves, estimate the cluster means from one half and score them against the other, so that the two share no counting noise. The cross-fitted oracle’s finest-band correlation is 0.031 [0.008, 0.134]. Means taken from the scored half itself give 0.031 [0.008, 0.134]. The shortfall ratio is 3.90 [2.23, 7.15] against 3.91 [2.23, 7.16]. Shared noise moves the median ratio by 0.002. The ratio exceeds one in 13 of 13 specimens. The learnable predictor of Table 2 uses cluster means from the training blocks. It gives the same picture without any test label: *β*_1_ = 0.04, a fine-band shortfall of 81% and a ratio of 3.96 [2.25, 7.56], above one in 13 of 13.

#### The image partition is a morphology partition, not a map of contiguous tissue areas

On the 8-neighbour grid graph a median 62% of edges join bins with the same label (range 44–69%), against 99% for the coordinate partition and 6% for a random one, but each cluster splits into a median of about 924 connected components, the largest holding 7% of its bins.

#### Symmetric and Gaussian operators leave the shortfall ratio in the same range as the lazy walk

Replacing the lazy random walk by its symmetrically normalised form (**I** + **D**^*−*1*/*2^**AD**^*−*1*/*2^)*/*2 gives a finest-band shortfall ratio of 3.70 [2.20, 7.33], and a Gaussian spatial kernel of width 16 *µ*m over neighbours within 48 *µ*m gives 3.90 [2.24, 7.35], against 4.04 [2.28, 7.69] for the default; each exceeds one in 13 of 13 specimens.

### K.5 Fifty-six further encoders under the same readout

Section 3.3 compares one trained model with the domain oracle. Table 18 repeats the whole computation for 57 encoders in all. Each uses the same ridge head, the same folds and the same operator. They span 6M to 1.1B parameters and cover general-purpose and pathology pre-training, including distilled students. The scalar compresses them. The 57 scores span 0.463 to 0.559, a range of 0.10. The finest-band correlations span 0.03475 to 0.23807, a factor of 6.9. Hibou-L leads on both the scalar and the finest band. The ranking is also scale-dependent. Against the scalar ranking, the finest band reorders 261 of the 1596 encoder pairs (Spearman 0.85), the *t* = 4 band 84 (Spearman 0.98) and the *t* = 32 band 132 (Spearman 0.96), so the same leaderboard reads differently at different widths. The scalar separates some pairs by less than 0.01. DINOv3-H+ and KEEP differ by 0.0039 in PCC and by 0.110 in *β*_1_. That is a 69% difference in finest-band fidelity relative to DINOv3-H+, and the scalar records it as a tie. The ranking is stable under resampling. We resample the 13 specimens with replacement (*B* = 2000). The finest-band ranking keeps a Spearman correlation of 0.97 [0.92, 1.00] with the full-data ranking, and the leader stays first in every resample. The direction holds for 138 of the 261 reversed pairs in at least 95% of resamples (189 at 80%). DINOv3-H+ leads KEEP in *β*_1_ by 0.068 to 0.124 across resamples, and the lead never changes sign. Per-encoder *β*_1_ intervals have a median width of 0.070. The result of Section 3.3 holds for all 57 encoders. Every encoder’s own domain oracle reaches 65–89% of its score. The finest-band shortfall exceeds the scalar shortfall by a factor of 2.4 to 5.9 across the 57.

**Table 18:** Fifty-seven encoders on the 22 Xenium regions under the same protocol. Rows are ordered by the encoder’s own score. Medians over specimens of per-specimen medians over regions; *β* at *t* = 32 has a median width of 52 *µ*m. M is the parameter count in millions, *β*_32_ the band at *t* = 32, Oracle/model the domain oracle’s share of the encoder’s own score, written as a fraction here and in per cent in the text, and Δ ratio the finest-band to scalar shortfall ratio of Section 3.3. The last entry of the right-hand block is the mean over the encoders.

| Encoder | M | PCC | $\beta_1$ | $\beta_{32}$ | Oracle/model | $\Delta$ ratio | Encoder | M | PCC | $\beta_1$ | $\beta_{32}$ | Oracle/model | $\Delta$ ratio |
| --- | --- | --- | --- | --- | --- | --- | --- | --- | --- | --- | --- | --- | --- |
| Hibou-L | 304 | 0.559 | 0.238 | 0.732 | 0.79 | 4.04 | LiteVirchow2 | 22 | 0.511 | 0.088 | 0.712 | 0.84 | 3.12 |
| Mascaret | 1136 | 0.552 | 0.224 | 0.734 | 0.81 | 4.13 | DistillPath-KS16 | 22 | 0.510 | 0.113 | 0.704 | 0.86 | 4.04 |
| Midnight-12k | 1136 | 0.550 | 0.228 | 0.736 | 0.78 | 3.40 | DistillPath-IS16 | 22 | 0.507 | 0.109 | 0.701 | 0.87 | 4.07 |
| Virchow | 631 | 0.548 | 0.197 | 0.735 | 0.81 | 3.72 | GigaPath-Flash | 22 | 0.506 | 0.126 | 0.702 | 0.88 | 3.71 |
| OpenMidnight | 1135 | 0.545 | 0.209 | 0.731 | 0.65 | 2.44 | Kaiko-L14 | 304 | 0.505 | 0.132 | 0.680 | 0.74 | 2.88 |
| H-optimus-1 | 1135 | 0.542 | 0.163 | 0.742 | 0.83 | 4.16 | Kaiko-S16 | 22 | 0.504 | 0.114 | 0.694 | 0.87 | 3.57 |
| H-optimus-0 | 1135 | 0.542 | 0.173 | 0.738 | 0.83 | 4.18 | Lunit-S16 | 22 | 0.501 | 0.136 | 0.687 | 0.89 | 5.88 |
| Virchow2 | 631 | 0.537 | 0.139 | 0.735 | 0.82 | 3.04 | DINOv3-H+ | 841 | 0.499 | 0.160 | 0.676 | 0.76 | 3.73 |
| Prov-GigaPath | 1135 | 0.537 | 0.182 | 0.724 | 0.83 | 3.93 | PathGen-CLIP | 428 | 0.498 | 0.123 | 0.684 | 0.86 | 4.39 |
| H0-mini | 86 | 0.536 | 0.165 | 0.729 | 0.85 | 4.66 | Kaiko-B16 | 86 | 0.498 | 0.117 | 0.691 | 0.83 | 3.61 |
| GPFM | 303 | 0.534 | 0.159 | 0.728 | 0.83 | 4.58 | CONCH | 395 | 0.496 | 0.147 | 0.679 | 0.84 | 4.46 |
| GenBio-PathFM | 1134 | 0.532 | 0.148 | 0.730 | 0.82 | 4.90 | KEEP | 414 | 0.496 | 0.050 | 0.700 | 0.84 | 4.22 |
| UNI v2 | 681 | 0.532 | 0.120 | 0.736 | 0.82 | 4.61 | Lunit-S8 | 22 | 0.496 | 0.073 | 0.690 | 0.88 | 5.36 |
| Phaet | 303 | 0.531 | 0.168 | 0.718 | 0.83 | 4.25 | CONCH v1.5 | 306 | 0.494 | 0.153 | 0.676 | 0.86 | 4.38 |
| LiteFM-L | 87 | 0.528 | 0.148 | 0.725 | 0.84 | 3.46 | CTransPath | 28 | 0.492 | 0.100 | 0.685 | 0.84 | 3.95 |
| Lunit-R50-BT | 24 | 0.527 | 0.168 | 0.709 | 0.85 | 4.38 | SigLIP 2 | 1136 | 0.486 | 0.111 | 0.670 | 0.81 | 3.80 |
| LiteFM | 22 | 0.524 | 0.138 | 0.721 | 0.84 | 3.41 | CIGA | 11 | 0.484 | 0.104 | 0.663 | 0.86 | 3.10 |
| Pathryoshka-B | 86 | 0.523 | 0.127 | 0.720 | 0.85 | 4.84 | RetCCL | 24 | 0.484 | 0.092 | 0.677 | 0.84 | 3.42 |
| Phikon-v2 | 303 | 0.521 | 0.134 | 0.719 | 0.83 | 4.25 | OmiCLIP | 638 | 0.480 | 0.104 | 0.661 | 0.82 | 4.04 |
| Hibou-B | 86 | 0.520 | 0.126 | 0.711 | 0.85 | 5.61 | DINOv3-L | 303 | 0.475 | 0.069 | 0.672 | 0.79 | 3.73 |
| Path Foundation | 27 | 0.520 | 0.143 | 0.706 | 0.86 | 4.57 | DINOv2-g | 1136 | 0.474 | 0.049 | 0.671 | 0.77 | 3.21 |
| mSTAR | 303 | 0.518 | 0.115 | 0.719 | 0.83 | 4.37 | DINOv3-B | 86 | 0.474 | 0.074 | 0.667 | 0.84 | 4.11 |
| UNI | 303 | 0.518 | 0.116 | 0.719 | 0.83 | 4.32 | PLIP | 151 | 0.471 | 0.038 | 0.669 | 0.86 | 3.82 |
| Lunit-R50-MoCo | 24 | 0.518 | 0.152 | 0.698 | 0.84 | 4.17 | CLIP-L/14 | 428 | 0.471 | 0.084 | 0.655 | 0.77 | 3.13 |
| Kaiko-B8 | 86 | 0.515 | 0.129 | 0.710 | 0.85 | 3.86 | DINOv2-L | 304 | 0.470 | 0.045 | 0.666 | 0.75 | 2.79 |
| Lunit-R50-SwAV | 24 | 0.514 | 0.145 | 0.703 | 0.85 | 3.36 | QuiltNet | 151 | 0.469 | 0.035 | 0.669 | 0.84 | 3.47 |
| Phikon | 86 | 0.514 | 0.107 | 0.713 | 0.84 | 4.31 | DINOv2-B | 87 | 0.465 | 0.045 | 0.658 | 0.77 | 2.56 |
| MUSK | 675 | 0.513 | 0.137 | 0.707 | 0.84 | 4.80 | BiomedCLIP | 196 | 0.463 | 0.097 | 0.637 | 0.89 | 4.72 |
| LiteFM-S | 6 | 0.512 | 0.114 | 0.712 | 0.84 | 3.25 | Mean over 57 | – | 0.509 | 0.126 | 0.701 | 0.83 | 3.97 |

### K.6 The benchmark’s finest band

Section 3.3 measures the finest-band shortfall on Xenium, where the bins are 16 *µ*m. The benchmark’s spots lie on a ∼ 100 *µ*m pitch. We repeat the same computation there. Table 19 gives the scalar and finest-band scores for all 57 encoders, using the same domain oracle and the same operator as Appendix C. The finest band on this pitch has a median width of 50 *µ*m, half the spot spacing. The domain oracle is above the ridge model on the scalar in 10 of 10 cohorts for every encoder but one. It stays above the model in the finest band as well, in a median 7 of 10 cohorts (range 3–9). Across encoders, 49 of the 57 have the oracle ahead in a majority of cohorts, and none have it ahead in all ten. Here the oracle leads at both scales, by a median 37% on the scalar and 15% in the finest band. Each lead is formed per sample and then reduced to its median within cohort, over cohorts and over encoders, so neither is the quotient of the levels in Table 19. At 100 *µ*m the finest accessible band is still coarser than the 10–30 *µ*m at which the model and the oracle separate on Xenium. So a predictor that knows only domain identity keeps a positive median band lead at the finest scale the benchmark can resolve, for 55 of the 57 encoders. The model of Section 3.3 recovers structure inside domains at 10–30 *µ*m, and the benchmark does not measure it, so no score on this pitch can directly validate that recovery.

**Table 19:** Scale-resolved comparison of the domain oracle and the ridge model on the benchmark. Each entry is formed per sample, medianed within cohort and then over cohorts, and summarised over encoders; the two cohort counts are per-encoder counts summarised the same way. Ridge uses the cohort-selected penalty of Section 3.2.

| Quantity | Median over 57 encoders | Range |
| --- | --- | --- |
| Scalar PCC, ridge | 0.335 | 0.252–0.387 |
| Scalar PCC, domain oracle | 0.482 | 0.353–0.505 |
| Finest-band $\beta_1$, ridge | 0.155 | 0.093–0.191 |
| Finest-band $\beta_1$, domain oracle | 0.169 | 0.112–0.195 |
| Cohorts with oracle above ridge, scalar | 10/10 | 5–10 |
| Cohorts with oracle above ridge, finest band | 7/10 | 3–9 |
| Finest-band width $\sigma_1$ ($\mu\text{m}$), set by the grid | 50.1 | 50.1–50.1 |

### K.7 An end-to-end super-resolution method at 16 *µ*m

iStar (Zhang et al., 2024) predicts expression at sub-spot resolution from H&E, through HIPT features and a small network. We ran it on the 16 Xenium regions of the first release under a matched protocol: within each region a spatial half split with 320 *µ*m blocks and a 64 *µ*m exclusion buffer, the 200 genes of Section 3.3 as its training targets, its own feature extraction and training on the training half, and its super-resolved output averaged over the 3 × 3 super-resolution cells around each held-out bin. Every predictor in Table 20 is scored on the same bins with the same operator. iStar reaches a median scalar score of 0.386 against 0.536 for ridge on Hibou-L features. Its finest-band correlation is 0.077 against 0.245. It is below ridge in 8 of 8 specimens for the scalar and 8 of 8 for the finest band. Against the domain oracle (0.410, 0.046) it is lower on the scalar and higher in the finest band. Its finest band holds 1.4% of its variance against 3.7% for ridge and 18.6% for the measurement, so the super-resolved field is smoother than the ridge prediction, not sharper. We also add spatial context to ridge: the mean features of the 8 neighbours and of the second ring within 48 *µ*m. This raises the scalar to 0.541 and the finest band to 0.269, a small gain concentrated in the finest band. To separate the backbone from the method we also fit ridge on iStar’s own HIPT features, sampled at the same bins on the same grid: it scores 0.429 on the scalar and 0.094 in the finest band (0.438 and 0.096 with *λ* = 10^4^), slightly above iStar itself and far below ridge on Hibou-L features. Most of the gap therefore comes from the backbone. The super-resolution head adds nothing over a linear readout of the same features. The finest band separates the two backbones by a factor of 2.5 at the same penalty. On the scalar the weaker backbone is only 18% behind, as a ratio of the two medians at the same penalty *λ* = 10^4^. This is one method under one protocol, but it shows that a method whose stated purpose is resolution does not automatically score higher where resolution is measured. The scalar shows the gap but understates its size. On the scalar iStar trails ridge by 25%. In the finest band it trails by 67%. Both are median paired shortfalls over specimens, taken against the median ridge score.

**Table 20:** An end-to-end super-resolution method against ridge on the Xenium regions, half split. Medians over specimens of per-specimen medians over regions, with the range over specimens; the last column is the finest band’s share of each field’s variance.

| Predictor (Xenium, half split, 200 genes) | PCC | $\beta_1$ | finest-band variance share |
| --- | --- | --- | --- |
| iStar (super-resolution, end-to-end) | 0.386 [0.242, 0.513] | 0.077 [0.025, 0.138] | 1.4% |
| Ridge on iStar’s own HIPT features (official $\lambda$) | 0.429 [0.295, 0.538] | 0.094 [0.038, 0.169] | – |
| Ridge on iStar’s own HIPT features ($\lambda = 10^4$) | 0.438 [0.303, 0.537] | 0.096 [0.040, 0.170] | – |
| Ridge on frozen Hibou-L features | 0.536 [0.404, 0.590] | 0.245 [0.114, 0.339] | 3.7% |
| Ridge with spatial context | 0.541 [0.409, 0.596] | 0.269 [0.134, 0.353] | 3.6% |
| Image partition, means from training blocks | 0.398 [0.297, 0.509] | 0.045 [0.013, 0.073] | 4.1% |
| Domain oracle | 0.410 [0.311, 0.515] | 0.046 [0.013, 0.077] | 4.1% |
| Measured field | – | – | 18.6% |

